# Structural determinants of microtubule minus end preference in CAMSAP CKK domains

**DOI:** 10.1101/586230

**Authors:** Joseph Atherton, Yanzhang Luo, Shengqi Xiang, Chao Yang, Kai Jiang, Marcel Stangier, Annapurna Vemu, Alexander D. Cook, Su Wang, Antonina Roll-Mecak, Michel O. Steinmetz, Anna Akhmanova, Marc Baldus, Carolyn A. Moores

## Abstract

CAMSAP/Patronins regulate microtubule minus-end dynamics. Their end specificity is mediated by their CKK domains, which we proposed recognise specific tubulin conformations found at minus ends. To critically test this idea, we compared the human CAMSAP1 CKK domain (HsCKK) with a CKK domain from *Naegleria gruberi* (NgCKK), which has lost minus-end specificity. Near-atomic cryo-electron microscopy structures of HsCKK- and NgCKK-microtubule complexes show that these CKK domains share the same protein fold, bind at the intradimer interprotofilament tubulin junction, but exhibit subtly different footprints on microtubules. Whereas NgCKK binding does not alter the microtubule architecture, HsCKK remodels its microtubule interaction site and changes the underlying polymer structure because the tubulin lattice conformation is not optimal for its binding. NMR experiments show that HsCKK is remarkably rigid, supporting this remodelling ability. Thus, in contrast to many MAPs, CKK domains can differentiate subtly specific tubulin conformations to enable microtubule minus-end recognition.

## Introduction

The involvement of the microtubule (MT) cytoskeleton in numerous processes in eukaryotic cells is enabled by the diverse and adaptable properties of individual MTs. MTs act as tracks for molecular motors, while growing and shrinking MTs can be used to generate force. MTs can also act as signalling hubs, such that specific tubulin conformations within particular regions of the polymer stimulate recruitment of distinct MT binding partners. The molecular basis of these effects, mediated by the conformational adaptability of tubulin dimers, is only just beginning to be understood and represents a key topic in the cytoskeleton field.

The ends of MTs are important sites of conformational diversity and are often points of communication between the MT cytoskeleton and other cellular components, such as membranes, organelles, centrosomes and chromosomes^1^. The exact conformation(s) of MT ends is an ongoing source of debate, but current evidence points to their being composed of zones with distinct and dynamic tubulin conformations ^2-7^. MT minus ends were long thought to be static and structurally homogeneous, capped by *γ*-TuRCs and buried at MT organizing centres. More recently however, the discovery and characterisation of CAMSAP (calmodulin-regulated spectrin-associated proteins)/Patronin family members has revealed that control of non-centrosomal MT minus-end dynamics, and their interaction with specific cellular regions, are vital in numerous aspects of cell physiology^8,9^. CAMSAP/Patronins are centrally involved in diverse activities including promoting cell polarity, regulation of neuronal differentiation and axonal regeneration, and definition of spindle organization and asymmetry, thereby highlighting the importance of regulation of MT minus-end dynamics in these varied contexts ^10-20^.

At the molecular level, CAMSAPs/Patronins stabilise uncapped MT minus ends and support MT minus-end growth^21-23^. CAMSAP/Patronins are large, multi-domain proteins with many cellular binding partners. However, the family is defined by the presence of a CKK domain (originally identified in <u>C</u>AMSAP1, <u>K</u>IAA1078 and <u>K</u>IAA1543) which is necessary and sufficient for MT minus-end binding in many CAMSAP/ Patronins^21,24^. Previously, we showed that CAMSAP/Patronin CKK domains preferentially bind to a zone behind the extreme MT minus end which corresponds to a region where the lattice undergoes a transition to gently curved tubulin sheets^2^. Subnanometer resolution single particle cryo-EM showed that CKK domains bind on the MT lattice between two tubulin dimers on adjacent protofilaments. Mutagenesis of residues at the MT-binding interface in the CKK domain disrupted lattice and minus end binding, showing that the same regions of the CKK domain that contact the MT are also involved in binding to the minus-end zone. Taking these data together, we proposed a model for CAMSAP/Patronin MT minus end recognition, which is mediated by sensitivity of the CKK domain to a curved sheet-like conformation of tubulin exclusive to MT minus ends. Specifically, the model suggests that the tighter CKK interaction with β-tubulin disfavours binding at MT plus ends while the looser α-tubulin contacts preferentially accommodate tubulin curvature at minus ends. This interaction also can occur on the MT lattice, but CKK binding induces distortion of the non-optimal binding site configuration, manifesting as protofilament skew within the polymer.

Despite these findings, several critical questions relating to the structural basis of this recognition mechanism remain unanswered: How is the CKK-induced MT lattice distortion accommodated, and what can this tell us about minus-end recognition? Can CKK binding to different MT protofilament architectures shed further light on the mechanism of minus end recognition? Intriguingly, we also previously identified CKKs in the amoeboflagellate *N. gruberi* and the potato blight fungus *P. infestans* that had lost the binding preference for MT minus ends that was proposed to be present in CKK domains from the last eukaryotic common ancestor^2^. Can a comparison of CKK domains with and without minus-end binding specificity also provide insight into MT minus-end recognition? Since discrimination between different MT lattice zones can depend on relatively subtle structural differences, high resolution information is needed to address these questions. It is currently not possible to image MT minus ends directly at the necessary resolution to observe these conformational variations. However, our previous work showed that CKK lattice binding can be used as a proxy for minus end binding, and cryo-EM studies of the lattice could yield near-atomic resolution information about the CKK-tubulin complex.

We therefore developed a new, RELION^25-27^-based image processing pipeline that enabled us to study the CKK-MT interaction at better than 4 Å resolution. We determined reconstructions of MT-bound CKK domains on both 13- and 14-protofilament pseudo-helical MTs, allowing visualisation of a wider repertoire of tubulin conformations. We also investigated the CKK domain from *Naegleria gruberi* (NgCKK), which does not show minus-end binding preference. The direct comparison of tubulin binding by NgCKK with that of human CAMSAP1 CKK (HsCKK) - which has minus end binding preference - allowed us to probe our previous model of minus-end recognition. We found that NgCKK, like HsCKK, binds between two tubulin dimers in neighbouring protofilaments. However, NgCKK MT binding is slightly shifted relative to HsCKK, resulting in a modified interaction with tubulin including smaller contacts with α-tubulin. NgCKK binding does not induce the protofilament skew, reinforcing that induction of skew is a structural signature for minus-end binding capability. The ability of HsCKK to remodel MTs in this way is supported by the distinctive rigidity of the HsCKK revealed at atomic resolution by NMR. HsCKK-induced skew arises within the MT lattice from the tilting of entire protofilaments coincidental with contraction of the MT diameter. These multi-disciplinary data reveal that the surprising structural plasticity of MTs, which is distinct from nucleotide-dependent modulations of the MT lattice, forms the basis for minus-end recognition by CAMSAP proteins.

## Results

### A CKK domain from *N. gruberi* shows no microtubule minus-end preference but binds at the canonical lattice CKK site

The presence of CKK domains in diverse organisms presents a unique resource that can shed light on the conserved or divergent properties of these domains. Our previous analysis suggested that a CKK domain from *N. gruberi* did not exhibit MT minus-end binding preference^2^. We confirmed this using TIRF experiments, showing that, in comparison to the well-characterised MT minus-end preference of HsCKK (Fig. 1a, left panel), NgCKK strongly bound along the entire MT lattice and exhibited no MT minus-end preference on dynamic MTs at a range of concentrations (Fig. 1a, right panels).

**Figure 1.**
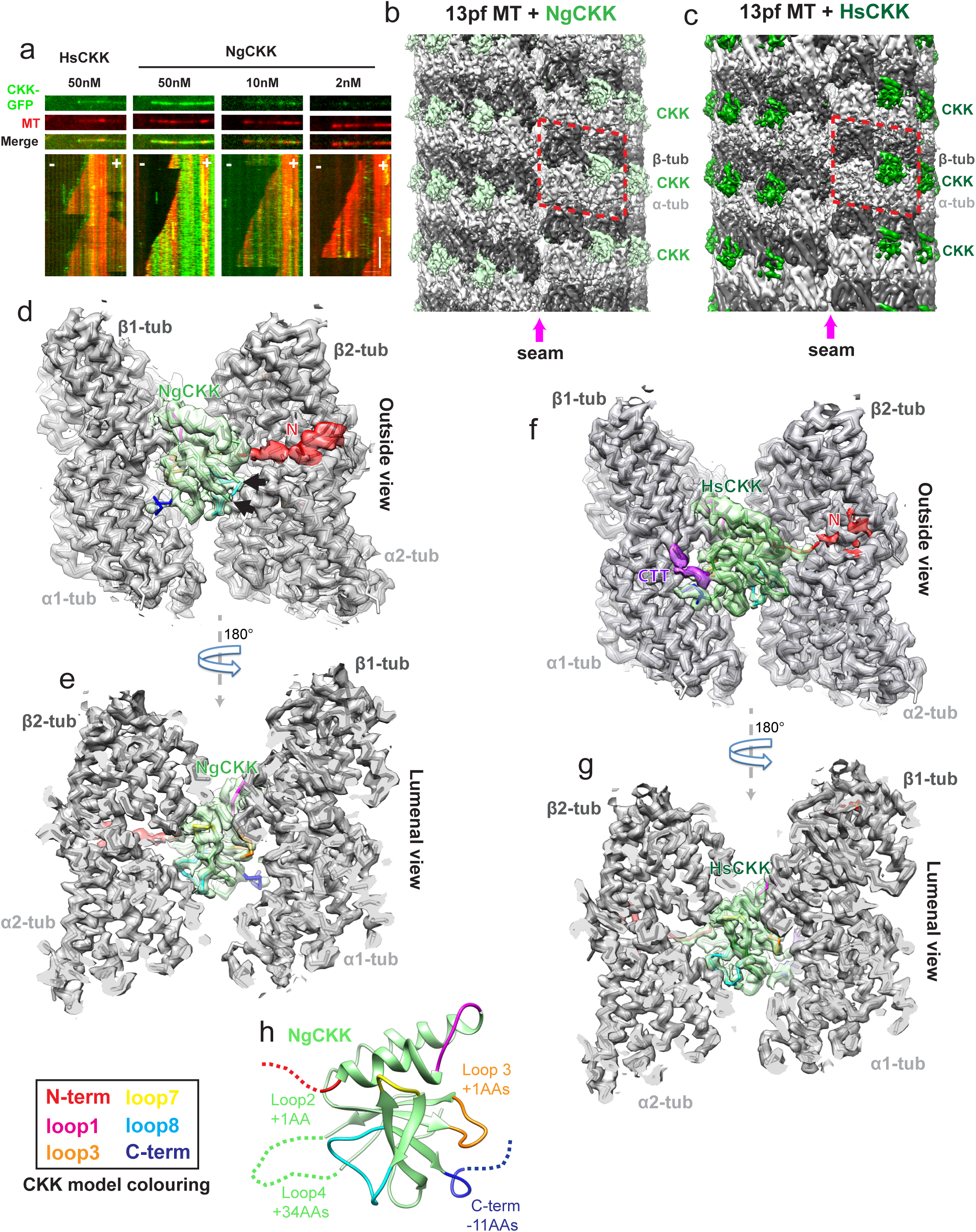
The CKK domain from *N. gruberi* does not recognise MT minus ends but, like HsCKK, binds at the intradimer interprotofilament binding site. a) TIRF assays with dynamic MTs demonstrate that unlike HsCKK, the NgCKK domain does not have a preference for MT minus ends but binds along the whole MT lattice at a range of concentrations. Scale bars: horizontal, 2 µm; vertical, 2 min. b) C1 reconstruction of the NgCKK-bound tsA201-tubulin 13-protofilament MT at 4.3Å resolution, showing CKK density (in light green) bound every 8nm between protofilaments at the intradimer tubulin interface, except at the MT seam; the reconstruction procedure produces a resolution gradient in the density which highest in the middle and lowest at the top and bottom; c) C1 reconstruction of the HsCKK-bound tsA201-tubulin 13-protofilament MT at 4.5Å resolution, showing CKK density (in green) bound every 8nm between protofilaments at the intra-dimer tubulin interface, except at the MT seam; as above, the reconstruction procedure produces a resolution gradient in the density which highest in the middle and lowest at the top and bottom; d) Asymmetric unit of the MT lattice extracted from the symmetrized 13-protofilament reconstruction viewed from the outside of the MT reveals the NgCKK-tubulin interaction, including ordering of the CKK N-terminus (red). The CKK model coloured according to CKK SSE/loop colour scheme at the bottom of the figure. Arrows show the start and end of the disordered 34 amino acid loop 4 insert in NgCKK; e) 180° rotated view of panel (d), showing the NgCKK interaction from the MT lumen with a cut through of the MT, showing the NgCKK as a wedge between 4 tubulin monomers. f) Asymmetric unit of the MT lattice extracted from the symmetrized 13-protofilament reconstruction viewed from the outside of the MT reveals the HsCKK-tubulin interaction, including partial ordering of the CKK N-terminus (red) and visualization of the unmodified tsA201 cell-tubulin β-tubulin C-terminal tail (purple). The CKK model coloured according to CKK SSE/loop colour scheme at the bottom of the figure; g) 180° rotated view of panel f, showing the CKK interaction from the MT lumen with a cut through the MT, showing the CKK as a wedge between 4 tubulin monomers and exhibiting well-ordered tubulin-interaction loops; h) NgCKK domain with loops coloured, indicating loop insertions and a shortened C-terminus (residue length differences in key loop regions compared to HsCKK are indicated).

To investigate this distinctive behaviour further, complexes formed by either NgCKK or HsCKK and taxol-stabilised MTs were imaged using cryo-EM for structure determination. Our previous work showed that CKK MT binding includes interactions with the C-terminal tails of tubulin (CTTs). To facilitate visualisation of this interaction, we used MTs assembled from tubulin purified from a human tsA201 cell line for our reconstructions^28^. These MTs contain only two β-tubulin isoforms and one α-tubulin isoform, have no detectable post-translational modifications as indicated by mass spectrometric analyses^28^, and are thus much more homogenous than the brain tubulin we previously used. To visualise CKK binding at higher resolution, we also developed a new image-processing pipeline for pseudo-helical MTs and different protofilament architectures in *RELION* (see methods). The resulting unsymmetrised (C1) reconstructions for both NgCKK and HsCKK showed distinct CKK intra-dimer, inter-protofilament densities every 8 nm along the MT axis and an absence of CKK density at the seam (Fig. 1b, c). This validates the accuracy of the pipeline and is consistent with our previous work^2^, while revealing the MT-bound NgCKK and HsCKK complexes at substantially higher resolutions now for both 13-and 14-protofilament MTs. The C1 reconstructions all have resolutions of 4.7Å or better, and the symmetrised reconstructions have resolutions of 3.8Å or better (Supplementary Fig. 1). This allowed us to build atomic models of the NgCKK-MT and HsCKK-MT complexes (Fig. 1d-g, Supplementary Table 1).

While the structures of mammalian CKK domains have previously been determined, our NgCKK reconstructions now reveal the near-atomic resolution structure of a non-mammalian CKK domain (Fig. 1h). It has a typical CKK fold, but sequence differences compared to HsCKK (Supplementary Fig. 2a) are reflected in structural differences in several loop regions (Supplementary Fig. 2bi). NgCKK’s loop4, which faces away from the MT surface (Fig. 1d, black arrows), is 34 amino acids longer than in HsCKK. There is no extra density in the cryo-EM reconstructions corresponding to this insert even at low thresholds (Fig. 1d), suggesting it is highly flexible/disordered and unlikely to be involved in MT binding. However, there are also structural differences in regions closer to the MT surface: specifically, loop3, the C-terminal single-turn helix and the beta-hairpin leading into loop7, all show backbone RMSDs >2.5Å (Supplementary Fig. 2bii).

The reconstructions show that both NgCKK and HsCKK form an intra-dimer inter-protofilament wedge (Fig. 1d-g), contacting both α-and β-tubulin subunits in a tubulin dimer pair (with constituent monomers numbered α1, β1, α2, and β2). Both CKK domains form MT contacts mainly via a set of surface exposed loops (Fig. 1d-g, described below). The N-terminal extension of each CKK domain also contacts the MT but in distinct ways. The NgCKK N-terminus forms an ordered density that is associated with the surface of β2-tubulin, although the density was not sufficiently defined to allow accurate modelling (Fig. 1d). Conversely, the HsCKK N-terminus closest to the CKK core forms a distinct interaction with β2-tubulin that was built into the atomic model, whereas density corresponding to its most N-terminal part was hardly visible on the surface of β2-tubulin (Fig. 1f). Furthermore, there is no clear contact between NgCKK and the β1-tubulin CTT (Fig. 1d). This is in contrast to HsCKK, where density corresponding to an additional ∼5 residues of β-tubulin’s CTT were visualised in our structures (Fig. 1f, purple) contacting both the CKK core and its C-terminus (Supplementary Fig. 3a). The visualisation of β-tubulin’s CTT interaction with HsCKK is likely facilitated by the limited sequence variability (Supplementary Fig. 3b) and lack of post-translational modifications in the CTT tails in tsA201 β-tubulin compared to brain tubulin^28^.

Overall, these structures show that the core of CKK domains with and without minus-end binding preference have the same protein fold and interact with MTs in similar ways. Differences in the tubulin interactions are seen at both their N-and C-termini, but our previous work showed that although these regions contribute to HsCKK MT affinity, neither the N-nor C-terminal extensions define its minus-end binding specificity. The differences in these regions are thus unlikely to explain the different end binding specificity of NgCKK and HsCKK. Instead, the observation of similar binding sites for NgCKK and HsCKK, together with our previous characterisation of HsCKK mutants, suggests that differences in CKK minus-end recognition behaviours are due to subtle structural differences in the interaction between tubulin and the CKK core. The near-atomic resolution of all our reconstructions allowed us to precisely investigate the mechanistic basis of these effects.

### Structural differences in MT interaction between NgCKK and HsCKK

To visualise how NgCKK’s MT interaction differs from HsCKK, each CKK-tubulin model was aligned on the tubulin parts of the complex, thereby revealing differences in CKK positioning relative to the MT lattice (Fig. 2ai). The tubulin dimers in both models readily superimpose (Supplementary Fig. 4), as do helix-α1, loop1 and loop7 in each CKK domain (Fig. 2ai). However, NgCKK loop3, loop8 and its C-terminus - which all contact the MT - are displaced compared to HsCKK (Fig. 2aii). Whilst some of these variations are due to structural divergence in NgCKK, this is insufficient to explain all the differences in CKK positioning relative to tubulin. In fact, NgCKK is rotated away from the MT around the apex of loop7 relative to HsCKK (Supplementary Movie 1).

**Figure 2.**
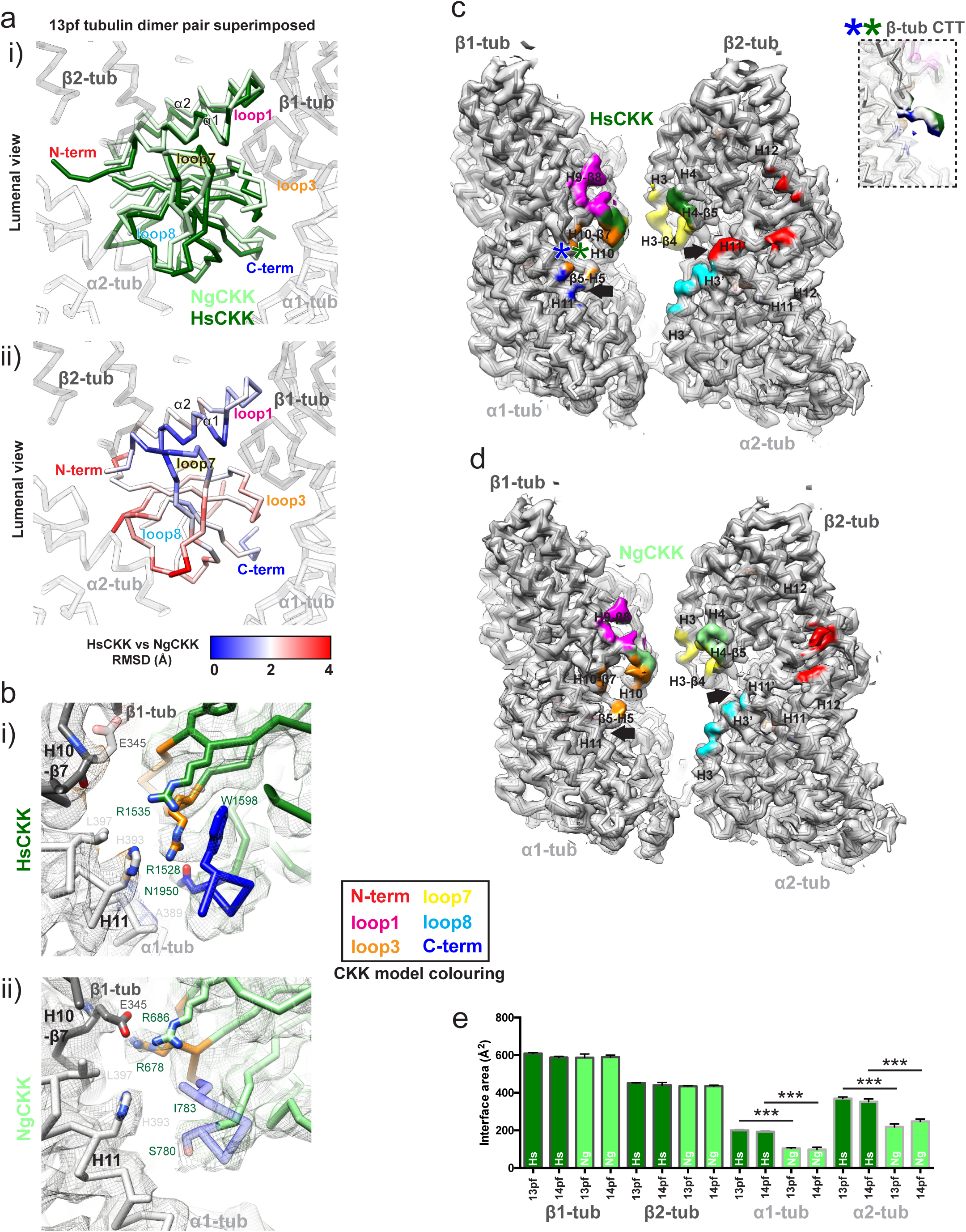
Comparison of MT-bound NgCKK and HsCKK structures reveal subtle differences in their domain structure and binding footprints. a) Differences in HsCKK and NgCKK positioning relative to tubulin (used for alignment and shown as white backbone model) viewed from the MT lumen; i) Overlay of HsCKK (dark green) and NgCKK (light green) backbones; ii) Backbone RMSD between HsCKK and NgCKK shown on the HsCKK structure, demonstrating their shifted binding position relative to tubulin particularly in tubulin-contacting loop8 and loop3. b) Comparison of the contacts formed between HsCKK or NgCKK and α-tubulin H11 (light grey backbone) and β-tubulin’s H10-β7 loop (dark grey backbone); i) HsCKK contacts with tubulin; ii) NgCKK contacts with tubulin; cryo-EM density is shown in mesh and the CKK model loops are coloured according to CKK SSE/loop colour scheme in the box. c) Overview of the HsCKK MT binding footprint on its intradimer interprotofilament binding site; tubulin density <5Å from HsCKK is coloured according to the CKK SSE/loop colour scheme in (b); asterisks refer to the inset, which shows HsCKK interaction with the β-tubulin CTT; arrows indicate regions where tubulin contacts differ in HsCKK and NgCKK. d) The footprint of NgCKK on the MT is different compared to HsCKK. Tubulin density <5Å from the CKK is coloured according to CKK SSE/loop colour scheme in (b). Arrows indicated regions where tubulin contacts differ in HsCKK and NgCKK. e) Calculated interface area in Å^2^ between HsCKK (dark green bars) or NgCKK (light green bars) and dimer pair subunits, α1-tubulin, α2-tubulin, β1-tubulin and β2-tubulin, for both 13- and 14-protofilament reconstructions, showing smaller α-tubulin contacts in NgCKK compared to HsCKK. Measurements were made in PISA using the final models that include tubulin contacts of 4 CKK domains (n=4; *** p<0.001, one-way ANOVA with Tukey’s multiple comparisons test).

As a result of this shift, even when the sequences in each CKK domain are conserved, our reconstructions show that some NgCKK residues engage differently with the MT lattice compared to equivalent residues in HsCKK. For example, HsCKK loop3 residues R1535 and R1528 contribute significantly to MT affinity. With the improved resolution of our current reconstructions, R1528 is observed extending close to α1-tubulin’s residues H393 on H11, whilst R1535 reaches to contact the β1-tubulin CTT (Fig. 2bi). On the other hand, in NgCKK loop3 – which is one residue longer and positioned differently with respect to the MT surface compared to HsCKK – the residue equivalent to R1528 (R678) interacts with loop H10-β7 of β1-tubulin, possibly hydrogen bonding with E345 (Fig. 2bii). In addition, instead of interacting with α1-tubulin, NgCKK residue R686 (equivalent to R1535) also interacts with β1-tubulin, extending to within hydrogen bonding distance of E345.

Altogether, the subtle differences within the NgCKK sequence, fold and positioning of the domain relative to tubulin result in a different binding footprint on the MT surface compared to HsCKK (Fig. 2c,d). The differences are most striking on α-tubulin, with the HsCKK C-terminus/loop3 and loop8/N-terminus forming closer contacts with α1-and α2-tubulin, respectively, compared to NgCKK. Indeed, the overall NgCKK footprint is significantly smaller on both α1-and α2-tubulin compared to HsCKK (Fig. 2e). This key observation supports the previously proposed importance of contacts with α-tubulins in mediating the minus end specificity of HsCKK.

### HsCKK binding is more sensitive than NgCKK to microtubule lateral curvature

In binding between two tubulin dimers, CKK domains are well placed to sense changes in inter-tubulin lateral curvature, which affects the distance and angle between adjacent protofilaments. Previously, we proposed that sensitivity to lateral curvature was an important facet of HsCKK binding minus-end specificity. Since MTs with different protofilament numbers exhibit different lateral curvature, comparison of CKK binding in our 13-and 14-protofilament reconstructions allowed us to investigate this effect. Superposition of a single tubulin dimer from each of the 13-and 14-protofilament atomic models for NgCKK and HsCKK reconstructions shows that the interprotofilament lateral angle in 14-compared to 13-protofilament MTs is ∼2° shallower (Fig. 3). In response, NgCKK is only slightly altered in its binding site on 14-protofilament MTs (Fig. 3a, Supplementary Movie 2), but HsCKK experiences a larger displacement out of the inter-protofilament cleft on 14-protofilament MTs (Fig. 3b, Supplementary Movie 3). Even though the HsCKK interface with α-tubulin remains larger than that of NgCKK on 14-protofilament MTs (Fig. 2e), the comparison between different MT architectures suggests that in the context of decreased lateral curvature, HsCKK is more prone to being squeezed out of its binding site than NgCKK. This is presumably because it binds deeper between protofilaments compared to NgCKK (Figure 3).

**Figure 3.**
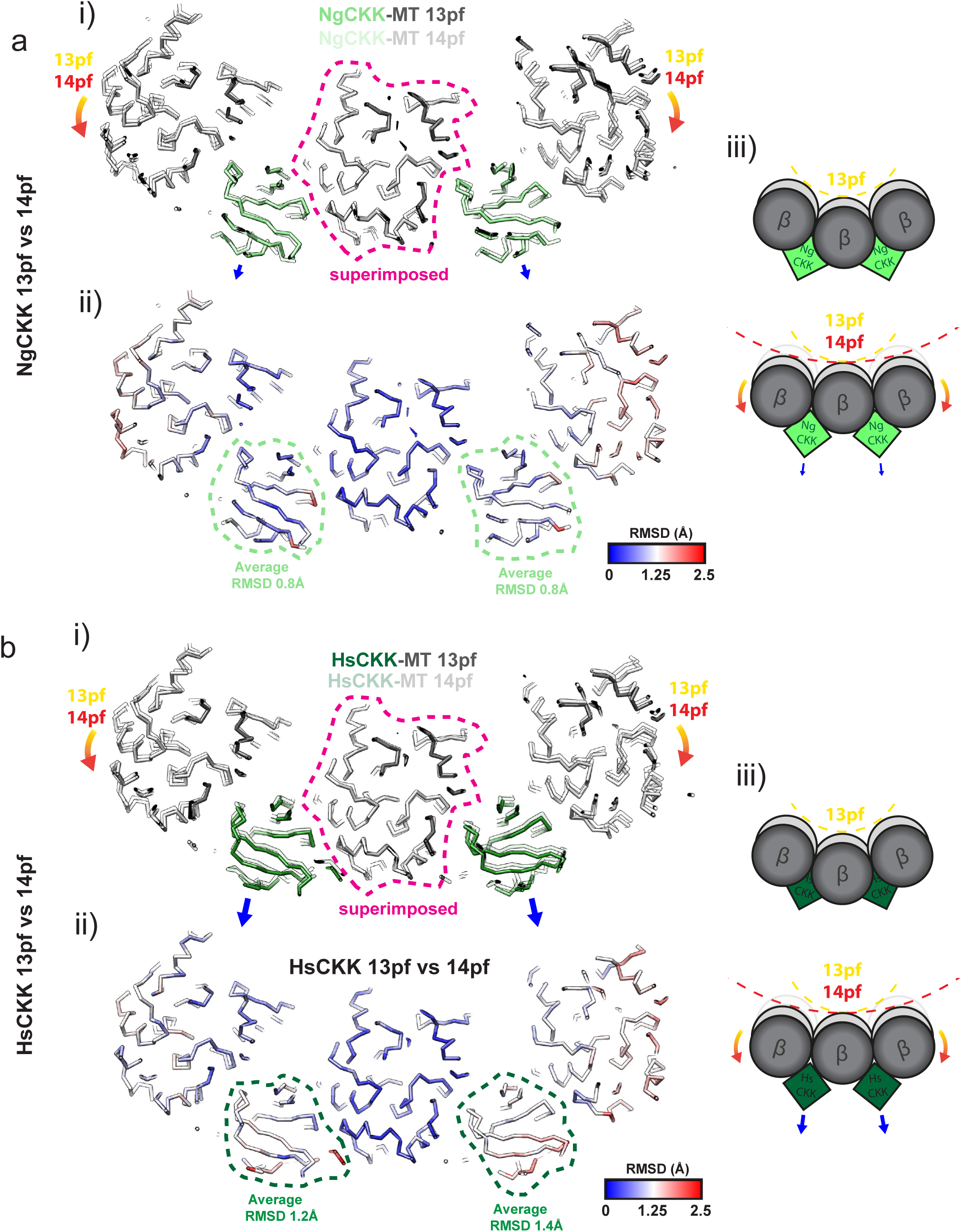
HsCKK is more displaced than NgCKK from its binding site on 14-protofilament compared to 13-protofilament MTs. a) A transverse slice, viewed from the MT plus end, through 3 protofilaments from the NgCKK-MT 13-and 14-protofilament models superimposed on the central protofilament. This shows that the adjacent protofilaments adopt a shallower relative angle in 14-protofilament MTs (orange arrows) and reveals the response of NgCKK bound in the interprotofilament valley to this change in lateral protofilament curvature. i) view of the superimposed protofilament backbone models from the 13-and 14-protofilament structures superimposed on the central protofilament (dotted pink outline); small blue arrows indicate the relatively small displacement of NgCKK from its binding site on 14-protofilament MTs. ii) Backbone RMSD between the 13-and 14-protofilament models shown on the 13-protofilament model; iii) schematic depicting the effect of MT protofilament architecture on NgCKK binding; b) A transverse slice, viewed from the MT plus end, through 3 protofilaments from the HsCKK-MT 13-and 14-protofilament models superimposed on the central protofilament. This again shows how the adjacent protofilaments adopt a shallower relatively angle in 14-protofilament MTs (orange arrows) and reveals the response of bound HsCKK to this change in lateral protofilament curvature. i) view of the superimposed protofilament backbone models from the 13-and 14-protofilament structures superimposed on the central protofilament (dotted pink outline); larger blue arrows indicate the larger displacement of HsCKK from its binding site on 14-protofilament MTs. ii) Backbone RMSD between the 13-and 14-protofilament models shown on the 13-protofilament model; iii) schematic depicting the effect of MT protofilament architecture on HsCKK binding;

Since our model of minus-end recognition predicts that lateral flattening of adjacent α-tubulin pairs would favour HsCKK binding, the outward displacement of HsCKK from its binding site on slightly flatter 14-protofilament MTs initially appears counter-intuitive. In the lattice, however, lateral flattening affects α-and β-tubulins equally, whereas our model suggested that both i) lateral flattening in α-tubulin towards the MT minus end is particularly important and furthermore that ii) preferential lateral flattening in adjacent β-tubulin pairs at MT plus ends would disfavour HsCKK binding. Thus, the potential enhancement of α-tubulin contacts in 14-protofilament MTs is balanced by tightening at the already tight β-tubulin interface. This emphasises the sensitivity to subtle differences in tubulin conformation encoded by HsCKK that supports its MT minus-end binding preference, in particular the importance of conformational asymmetry in α-and β-tubulins at this site, as we previously predicted.

### Ability of CKKs to induce protofilament skew correlates with minus end specificity and arises from tilting of intact protofilaments

In addition to comparison of the NgCKK and HsCKK binding sites, the overall architecture of the decorated MTs can be compared to shed further light on HsCKK minus-end preference. We previously described the ability of HsCKK to induce positive (right-handed) protofilament skew in MTs polymerized from mammalian brain tubulin. Raw images (Fig. 4a) and particle alignment parameters (Fig. 4b) from our new HsCKK-MT data set support this observation on tsA201 cell tubulin MTs. 13-protofilament MTs usually have unskewed protofilaments, running straight along the MT wall (Fig. 4c), whereas HsCKK binding causes right-handed protofilament skew (Fig. 4a-c). Furthermore, 14-protofilament MTs usually have negatively skewed protofilaments but HsCKK binding caused these protofilaments to lie parallel to the MT wall with no skew (Fig. 4a,b). In other words, we observed induction of right-handed protofilament skew in both types of MT architectures. Intriguingly -and in contrast to the HsCKK -the intrinsic protofilament skew in both 13-and 14-protofilament MTs was unperturbed by NgCKK binding (Fig. 4a-c). This is consistent with the idea that protofilament skew induction correlates with MT minus-end specificity.

**Figure 4.**
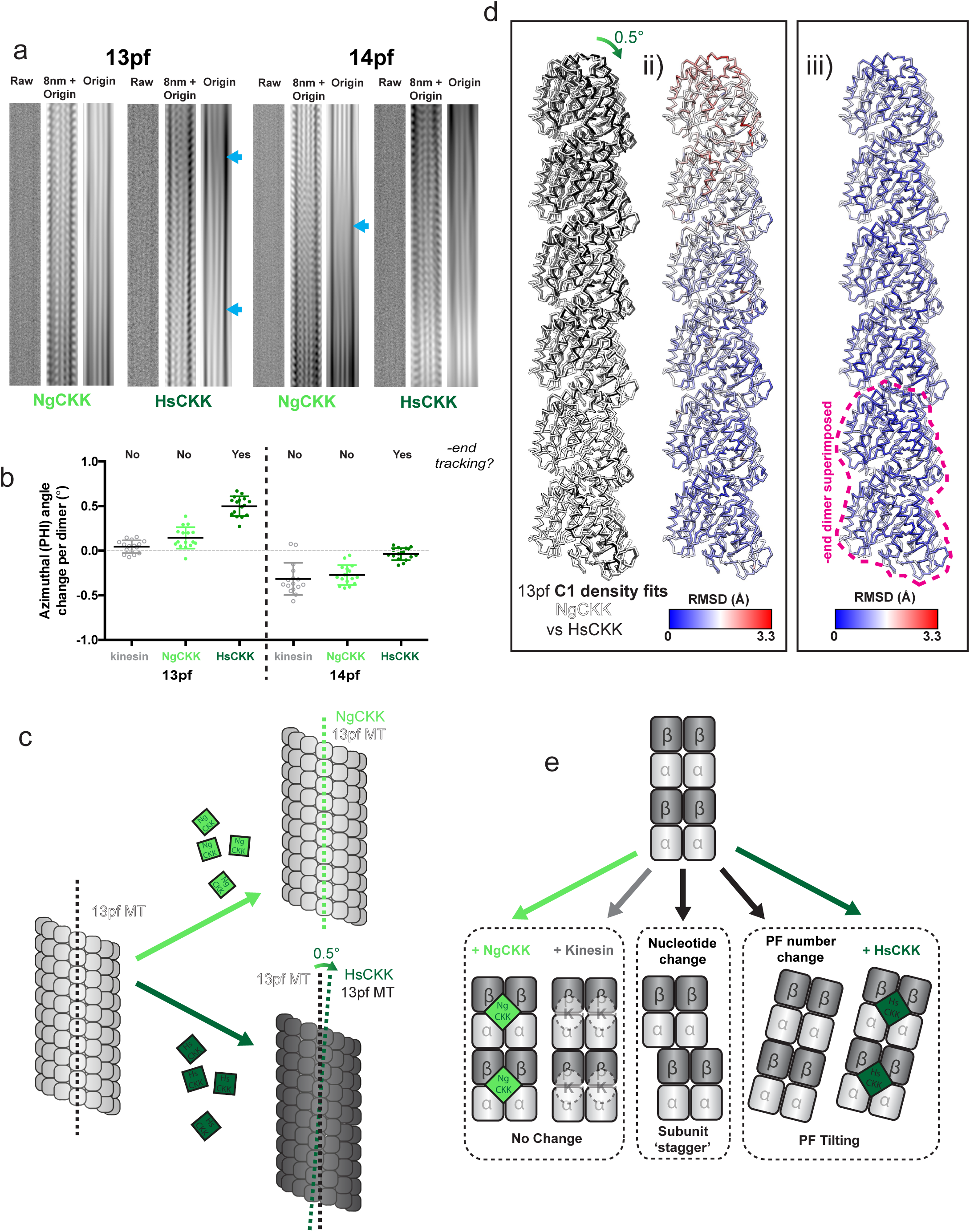
HsCKK, but not NgCKK, induces protofilament skew by tilting of whole protofilaments relative to MT axis. a) Raw and Fourier filtered images of 13-(left) and 14-protofilament (right) MTs decorated by NgCKK and HsCKK. In each set of 3 panels: left, raw image; centre, filtered image to include data at, and adjacent to, the origin and the 1/8nm layer line shows density corresponding to the CKK domain every tubulin dimer; right, filtered image to include data at, and adjacent to, the origin highlights the MT moiré pattern and the effect of CKK binding on protofilament skew; blue arrows indicate variations in the moiré pattern that arise from protofilament skew. b) Protofilament skew for a 16 MT subset from each dataset is depicted by plotting the average rotation angle around the MT axis (PHI) change per dimer moving axially towards the MT plus-end. 13-and 14-protofilament HsCKK-decorated MTs are compared to control kinesin decorated paclitaxel-stabilized MTs (13-protofilament kinesin-3 data from [ref]; 14-protofilament kinesin-1 data from [ref]) that show no skew for 13-protofilament and a left-handed (negative) skew for 14-protofilament MTs. Data represent mean ± SD. HsCKK 13-protofilament (tSA201 tubulin) vs kinesin-3 13-protofilament, p<0.0001, NgCKK 13-protofilament vs HsCKK 13-protofilament, p<0.0001, NgCKK 13-protofilament vs kinesin-3 13-protofilament, not significant (p=0.164), HsCKK 14-protofilament (tSA201 tubulin) vs kinesin-1 14-protofilament, p<0.001, NgCKK 14-protofilament vs HsCKK 14-protofilament, p<0.0001, NgCKK 14-protofilament vs kinesin-1 14-protofilament, not significant (p=0.889), one-way ANOVA with Tukey’s multiple comparisons test. c) Schematic of how protofilament skew in 13-protofilament MTs is influenced by NgCKK vs HsCKK binding d) i) MT protofilaments fitted into aligned HsCKK and NgCKK C1 reconstructions were overlaid, revealing the protofilament skew induced by HsCKK binding; the bottom dimer corresponds to the point at which the density was aligned; divergence between NgCKK and HsCKK models increases from this point; ii) RMSD of backbone positions in panel i) depicted on a NgCKK protofilament; iii) For comparison, RMSDs between NgCKK and HsCKK protofilaments calculated when the bottom dimers in the models themselves are directly aligned. RMSD does not decrease significantly with distance from the superimposed dimer, suggesting skew change introduced by HsCKK relates to tilt of whole protofilaments rather than stagger between dimers along a protofilament. e) schematic depicting mechanisms of MT protofilament skew induction: left, no skew change observed due to binding by NgCKK and kinesin motor domain; middle, protofilament skew arising from interdimer subunit stagger, e.g. arising from tubulin GTPase; right, protofilament skew arising from whole protofilament tilting relative to the MT axis e.g. due to changes in protofilament number or from binding by HsCKK.

Protofilament skew can arise either from a stagger of individual subunits along a protofilament perpendicular to the long axis of the protofilament, or from tilt of the whole protofilament relative to the MT axis^29^. The higher resolution of our new reconstructions allowed us to probe how protofilament skew is structurally accommodated in HsCKK-bound MTs and thereby shed light on requirements for tubulin plasticity to support HsCKK minus end recognition. To do this, we aligned and compared the HsCKK and NgCKK C1 cryo-EM reconstructions; this is because although these structures have slightly lower resolutions than the symmetrised reconstructions, the fact that they have not been symmetrised means they more closely reflect the overall polymer structure. When protofilaments of NgCKK and HsCKK models fitted into their corresponding C1 reconstructions are then overlaid, a positive skew of HsCKK protofilaments relative to NgCKK protofilaments is observed (clockwise rotation viewed from the outer surface the MT, Fig. 4di). This skew is reflected in increasing RMSD between the two models along the helical axis (shown in Fig. 4dii). However, when the models of individual protofilament from NgCKK and HsCKK structures are directly aligned, this produces only a small RMSD (<1Å) along the whole protofilament (Fig. 4diii), i.e. the structures of protofilaments from each reconstruction are essentially the same. This comparison shows that, rather than rearrangements within protofilaments, protofilament skew in HsCKK-bound MTs results from a tilt of whole protofilaments relative to the pseudo-helical axis (Fig. 4e).

### CKK domains are remarkably rigid both in the free and MT-bound state

What are the properties of HsCKK that support induction of whole-protofilament skew? To answer this question at atomic resolution we turned to solid-state NMR (ssNMR). To collect the highest quality ssNMR data, a high affinity HsCKK-MT interaction is required. Our previous work identified an HsCKK mutant, N1492A, that increased the binding affinity for MTs compared to wild-type HsCKK while reducing its selectivity for MT minus ends. Therefore, a range of NMR data were collected using this mutant, and in line with its previously described MT-binding properties, we observed improved resolution in our ssNMR spectra compared to the wild type CKK^2^ (Supplementary Fig. 5a). Importantly, this gain in spectral resolution allowed us to conduct fast magic angle spinning (MAS), ^1^H detected ssNMR experiments to further elucidate the CKK conformations in complex with MTs (Supplementary Fig. 5b and c). The chemical-shift perturbations observed upon MT binding agreed with the previous reported CKK-MT interface while providing a more quantitative and residue-specific description of the behaviour of the domain (Fig. 5ai).

**Figure 5.**
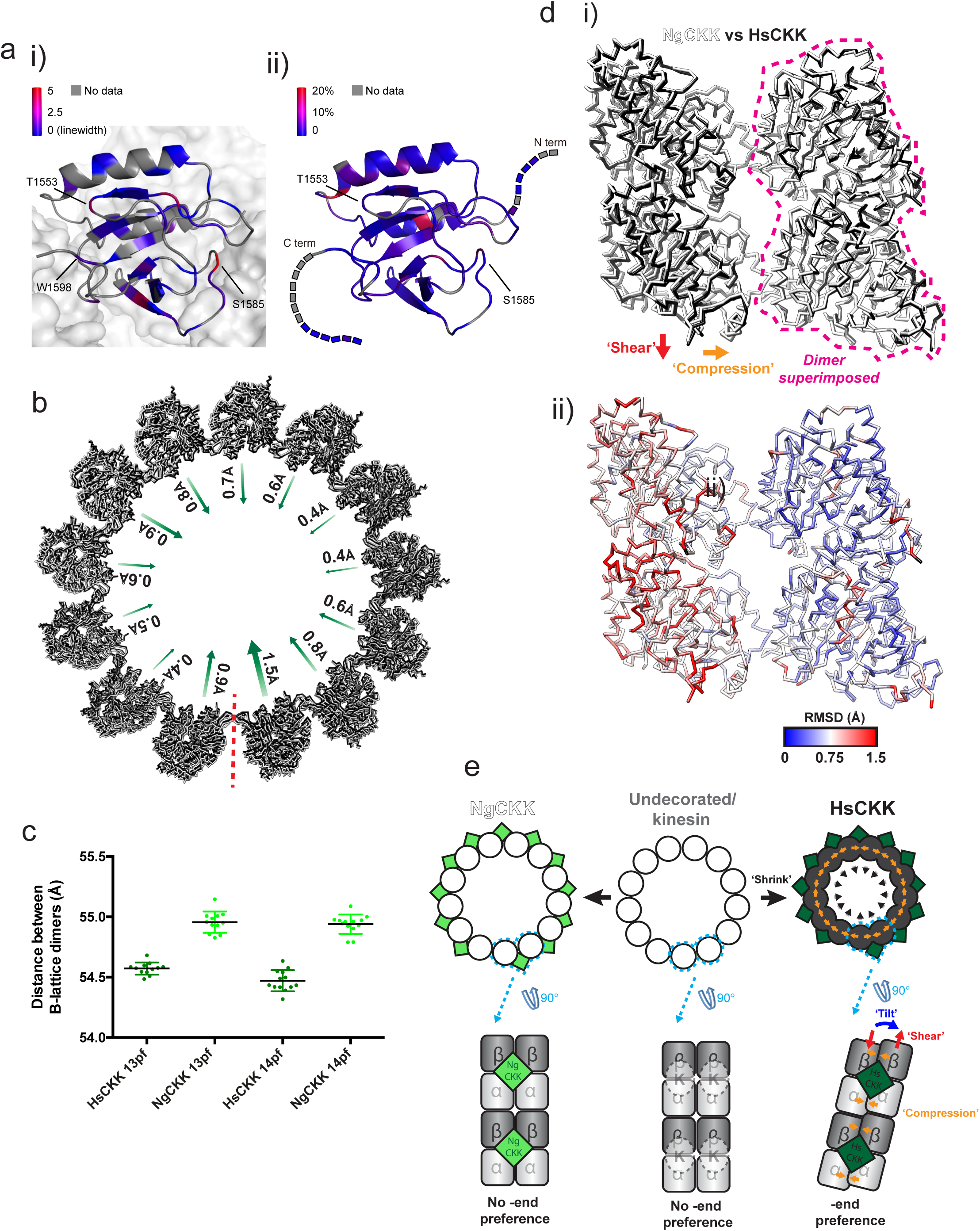
Rigidity of the HsCKK domain revealed by high-resolution NMR mediates local remodelling of the CKK MT binding site, and is accommodated by MT diameter shrinkage and protofilament shear and skew. ai) Chemical-shift perturbations, coloured by strength, arising from HsCKK_N1492A binding to MTs mapped on the structure of CKK-MT complex (PDB: 5M5C); the CKK domain is depicted in a ribbon representation while the MT surface is shown as a space-filling model; aii) Changes of transverse relaxation rates obtained from solution-state NMR CPMG experiments plotted on the 3D structure of the free HsCKK domain; b) HsCKK MT binding is accompanied by contraction of the MT diameter; a single turn of models docked within the aligned HsCKK and NgCKK C1 reconstructions are shown viewed from the minus end; arrows indicate the irregular shift of individual protofilaments; c) Contraction of MT diameter is caused by shrinkage of the distance between adjacent dimers; the distance between the centre of masses of each pair of adjacent B-lattice dimers was measured in 13-and 14-protofilament C1 HsCKK and NgCKK models; differences between HsCKK and NgCKK models are statistically significant (p<0.0001, t-test); di) Alignment of a single dimer from the NgCKK-tubulin and HsCKK-tubulin C1 models shows HsCKK induces compression and shear between the dimers at its binding site; ii) RMSD of backbone positions in panel i); e) Schematic summarising modifications imposed by HsCKK but not kinesin or NgCKK binding on MT architecture; modifications are exaggerated for clarity.

To obtain additional insight into the binding mechanism of CKK domains to MTs, we probed residue-specific dynamics of the free CKK domains using solution-state NMR. Specifically, we conducted Carr-Purcell-Meiboom-Gill (CPMG) relaxation dispersion^30,31^ and Chemical Exchange Saturation Transfer (CEST) experiments ^32^ to reveal possible millisecond time-scale conformational exchange processes of mammalian CKK domains. The CPMG profiles of HsCKK_N1492A, HsCKK and the CKK domain from mouse CAMSAP3 all showed no exchange in these CPMG time ranges (50 Hz∼1.5 kHz) (Fig. 5aii and Supplementary Fig. 5d). Similarly, the results of additional CEST experiments using HsCKK_N1492A speak against slow milli-second time-scale motion in free CKK domains, for example, in residues T1553 and S1585, which exhibited significant chemical-shift changes upon complex formation (Supplementary Fig. 5e). Similar results were obtained for CAMSAP3 CKK (Supplementary Fig. 5f). Taken together, our NMR experiments suggest that the 3D structures of human CKK domains are remarkably rigid and, in contrast to many other MAPs, do not undergo structural changes upon MT binding. These CKK properties are likely to be integral to the mechanism of CAMSAP CKK recognition of MT minus ends.

### HsCKK remodels its binding site causing protofilament tilting and MT diameter shrinkage

Given the unusual rigidity of HsCKK, we wanted to know how protofilament skew was brought about by HsCKK binding and accommodated by the MT lattice. Given the conservation of the longitudinal and lateral tubulin contacts, the geometric constraints that define MT architecture are captured in the lattice accommodation model^33,34^; we investigated in turn the set of interconnected structural parameters it describes.

The helical rise and monomer repeat distances were not significantly different in MTs (13-or 14-protofilament) bound by HsCKK or NgCKK (Supplementary Table 2). However, protofilaments in HsCKK-MTs are closer together (Fig. 5b), giving these MTs a ∼4 Å smaller diameter than NgCKK MTs with the same protofilament number. The diameter shrinkage occurs because the centres of mass of all neighbouring B-lattice dimers in a single 3-start helical turn are closer by around 0.4Å in HsCKK-compared to NgCKK-MTs (Fig. 5c). This inward protofilament positioning is not symmetrical around the MT, with a range of 0.4-1.5Å relative shifts observed in both 13-and 14-protofilament architectures (Fig. 5c), and with the biggest deviations seen at and opposite of the seam. The small compression between lateral neighbouring tubulin dimers in the lattice can be observed by superimposing the ones from the atomic models of NgCKK-MT and HsCKK-MT (Fig. 5d). This analysis also reveals a longitudinal displacement of adjacent tubulin dimers in the HsCKK model relative to the NgCKK model; this is indicative of shearing of adjacent protofilaments as they skew. There are, however, no detectable differences in inter-protofilament lateral contacts (small RMSDs, Fig. 5dii; Supplementary Fig. 6). Rather small adjustments across the outer tubulin surface -where HsCKK binds -flexibly accommodate shifts in the MT architecture due to HsCKK binding (Fig. 5dii, larger RMSDs in the tubulin on the left). In summary, relative to NgCKK, HsCKK induces small conformational changes in tubulin at its binding site which, in the context of whole MTs, induces protofilament tilt, shear, lateral compression and a reduction in MT diameter (Fig. 5e).

## Discussion

To shed light on the MT minus-end binding preference of CAMSAPs, we have structurally compared a CKK domain that does not bind MT minus ends – NgCKK – with the CKK domain from human CAMSAP1 (HsCKK), which mediates CAMSAP1’s MT minus-end binding preference. Little is known about the native MT ultrastructure of *Naegleria*, so it is possible that NgCKK could recognise MT minus ends on *Naegleria* MTs^35^. However, for the purposes of our current study, NgCKK has proven an invaluable tool for evaluating MT minus end binding mechanisms on mammalian MTs. To allow a near-atomic resolution investigation of the subtle mechanism(s) at work, we studied NgCKK and HsCKK MT lattice binding in the context of different MT architectures and used these structures to explain the differences in their MT minus-end recognition properties (Fig. 6).

**Figure 6.**
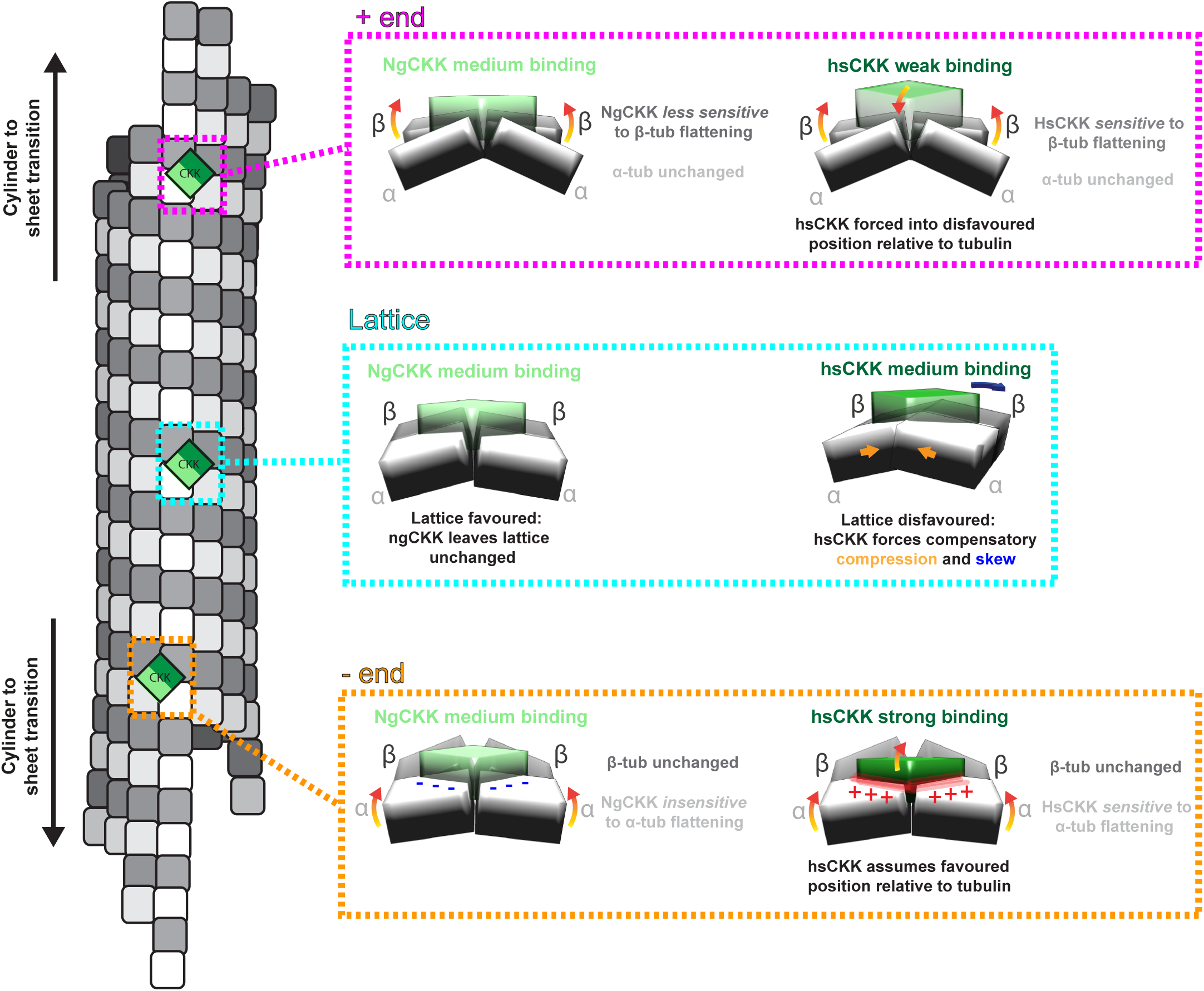
Comparison of MT binding properties of NgCKK and HsCKK substantiates and expands the model of CAMSAP MT minus end recognition. The schematic of a stable/growing 13-protofilament MT on the left shows different zones within the polymer in which the tubulins adopt subtly different conformations. Towards MT ends, there is a region of transition from the cylindrical lattice to curved sheet-like regions in which interprotofilament connections are maintained but which exhibit decreasing lateral curvature and increasing longitudinal curvature away from the MT lattice. Beyond this transition zone, protofilaments gradually terminate and separate at each MT end. On the right, the interaction of NgCKK and HsCKK with the unique tubulin dimer pair conformation in each zone is compared. On the MT lattice (middle, cyan boxed zone), as visualised in our cryo-EM reconstructions, NgCKK and HsCKK bind at the same site, but HsCKK remodels its binding site by compressing the tubulin dimer pairs, inducing protofilament skew. At the plus end lattice-end transition (top, pink boxed zone), we hypothesise that NgCKK is insensitive to the plus end specific tubulin sheet curvature found here, whereas HsCKK binding is actively disfavoured because of the enhanced lateral flattening in the β-tubulin pair that is specific to the MT plus end. At the minus end lattice-end transition (bottom, orange boxed zone), we again hypothesise that NgCKK is insensitive to the minus end specific sheet curvature, whereas HsCKK binding is favoured because the specific asymmetrically curved configuration of the tubulin pair – in particular the enhanced lateral flattening in the α-tubulin pair -is the preferred binding conformation for HsCKK, and in which contacts with the α-tubulin pair are enhanced.

We found that NgCKK and HsCKK share the same protein fold and they bind at the same intra-dimer interprotofilament site. This shows that the presence/absence of MT minus-end recognition is not due to large conformational changes in the domain or shifts in CKK binding site. However, HsCKK forms a more extended interface with the α-tubulin pair at this site than NgCKK and sits deeper within the inter-protofilament groove. We found that HsCKK is more displaced from its binding site on 14-compared to 13-protofilament MTs, squeezed outwards by the flatter lateral curvature of adjacent protofilaments in the higher protofilament number MT architecture. In addition, HsCKK binding induces positive protofilament skew in both 13-and 14-protofilament MTs while NgCKK does not. Our reconstructions show that HsCKK brings the tubulin dimers to which it binds closer together, while a combination of NMR methodologies reveal that HsCKK is remarkably rigid, supporting its ability to remodel its binding site. While neither intra-nor inter-dimer longitudinal interfaces are perturbed in HsCKK-bound MTs, their global lattice architecture alters to accommodate the local remodelling at the HsCKK binding site: compared to NgCKK-MTs, the diameter of HsCKK-MTs are smaller and adjacent protofilaments exhibit compression, shear and skew while conserving the B-lattice MT architecture with a single seam predicted by the lattice accommodation model.

The first new and important aspect of the MT minus-end recognition mechanism by HsCKK revealed by our current data is that the CKK domain itself does not flexibly respond to different tubulin conformations. Rather, its rigidity is consistent with its sensitivity to, and affinity for, the conformation(s) of polymerised tubulin it encounters. Second, we confirmed that the ability to induce significant protofilament skew in fully decorated MTs correlates with MT minus-end recognition activity. This was previously observed in the HsCKK-N1492A mutant but is now confirmed in the comparison of HsCKK with NgCKK. Third, we previously speculated that skew induction reflects the non-optimal geometry for HsCKK binding of tubulin dimers within the MT lattice compared to minus ends. Our new reconstructions show that this is indeed the case, and that skew arises in response to HsCKK forcing the two tubulin dimers it contacts closer together. Consistent with this idea, the CKK binding site on neighbouring tubulin dimers are predicted to be laterally closer in the transition zone to gently curved tubulin sheets near MT minus ends which HsCKK prefers. Fourth, a key prediction of our model is that end specificity by HsCKK is mediated by the asymmetric curvature of tubulin at the minus end, with the α-tubulins less laterally curved relative to the β-tubulins. Conversely, at plus ends -with the β-tubulins less laterally curved -HsCKK binding is inhibited. We observed more extensive contacts with α-tubulins by HsCKK compared to NgCKK that are likely to be important in this discrimination mechanism. Altogether, this model describes how HsCKK has highest affinity for MT minus ends, some capacity to interact with the MT lattice but lowest affinity for plus ends (Fig. 6). It is clear that small differences between HsCKK and NgCKK, and between the effects they induce on MT binding, combine to produce large effects in terms of MT end recognition properties.

Structural studies of MT-bound MAPs typically reveal conformational changes in the MAP on interaction with the MT lattice. In the most extreme cases, unstructured proteins such as members of the tau/MAP2 family and the mitotic regulatory protein TPX2, become at least partially ordered when in contact with MTs^36,37^. A recent study of the plant MAP Companion of Cellulose synthase 1 also showed similar behaviour in its disordered N-terminus on binding MTs^38^. Folded MT binding domains in a number of MAPs – for example, kinesin motor domains^39,40^, CH domains in EB proteins^41^, the p150glued CAP-Gly domain^42^ -often undergo some rearrangements and/or ordering of otherwise disordered loop regions on formation of the MT-bound complex. In contrast, we show that the core of HsCKK, which is essential for minus end recognition, is sufficiently rigid that it does not undergo conformational changes on MT interaction, but rather the MT lattice is remodelled in response to HsCKK binding. In the case of HsCKK, this is because the main MT lattice is not the preferred binding substrate for CKK. However, this behaviour - in which a structurally invariant MAP is exquisitely sensitive to the precise conformation of the underlying tubulin - is likely to be shared by other proteins.

The availability of increasing numbers of MT structures bound by a range of ligands has also emphasised that, far from being an inflexible, structurally invariant cylinder, the MT lattice supports surprising structural plasticity. Lattice compaction at the interdimer interprotofilament tubulin contacts in response to the tubulin GTPase is well documented in MTs polymerised from mammalian tubulin^29,41,43-45^. End Binding (EB) proteins bind at the corner of four tubulin dimers adjacent to the tubulin GTPase site^46^ and their preference for the sleeve of GDP.Pi compacted tubulins that dynamically evolves as MTs grow mediates their tip-tracking activity^41,46-48^. EB binding itself induces a small left-handed protofilament skew by introducing a slight interdimer stagger along the protofilament, the mechanistic significance of which is not yet understood. CKKs bind MTs 4 nm away from the EB binding site and are insensitive to nucleotide-dependent conformational changes in the lattice^2^. We show here that HsCKK binding induces right-handed protofilament skew via tilt and shear of entire protofilaments, a completely different mechanism than seen for EBs. Thus, our characterization of the HsCKK-MT interaction also highlights the extent of structural plasticity that can be accommodated in the MT lattice. Small conformational effects induced by one MAP could have substantial consequences for binding of other MT-binding factors. We previously demonstrated that CKK binding at MT ends can sterically compete with kinesin-13 at MT minus ends^2^, thereby protecting them from depolymerisation^21-23^. Our current work also suggests that, beyond direct steric competition, different MAPs may exert allosteric control over each other’s MT binding by modifying the conformation of the MT lattice^29^.

Taken together, our data support the idea that MTs can act as allosteric signalling platforms, in which the precise configuration of polymerised tubulins are influenced by their dynamic state and binding partners ^6,49^. In the case of CAMSAPs/Patronins, sensitivity to structural variations in tubulin is essential for MT minus end recognition. These insights will inform future mechanistic investigations of conformational signalling arising from the MT cytoskeleton.

## Acknowledgements

This work was funded by grants from the Medical Research Council, U.K. (MR/R000352/1) to C.A.M, from the Dutch Science Foundation NWO ((VENI grant 722.016.002 to S.X., NWO-Groot (no. 175.010.2009.002) and TOP-PUNT (no. 718.015.001) grant to M.B.) and by uNMR-NL, National Roadmap Large-Scale NMR Facility of the Netherlands (grant 184.032.207), supported by a China Scholarship Council scholarship to C.Y., the intramural program of the National Institute of Neurological Disorder and Stroke (NINDS) and National Heart, Lung and Blood Institute (NHLBI) for A.R.M., and funding from the Swiss National Science Foundation (31003A_166608) to M.O.S..

## Author contributions

J.A., Y.L., A.A., M.B., C.A.M. designed the research. M.S., A.V., A.C., S.W., A. R-M., M.O.S., supplied reagents and analytical methods. J.A., Y.L., S.X., C.Y., M.S. performed the research. J.A., Y.L., S.X., C.Y., K.J., M.S., A.A., M.B., C.A.M. analysed the data, and J.A., Y.L., M.B., and C.A.M. wrote the manuscript, with contributions from all authors.

## Materials and Methods

### Protein expression and purification for TIRF Microscopy

Strep-GFP-tagged human CAMSAP1 CKK (residues 1474-C) and Naegleria gruberi CKK (residues 612-C, reference sequence XM_002675733.1) were prepared as described previously (Atherton et al., 2017). Briefly, both proteins were expressed in HEK293T cells using a modified pTT5 expression vector (Addgene no. 44006), purified using StrepTactin beads (GE) and eluted in elution buffer (50 mM HEPES, 150 mM NaCl, 1mM MgCl_2_, 1mM EGTA, 1mM dithiothreitol (DTT), 2.5 mM d-Desthiobiotin and 0.05% Triton X-100, pH 7.4). Purified proteins were snap-frozen and stored at −80 °C.

### TIRF microscopy analysis of CKK binding to dynamic MTs

TIRF microscopy was performed on an inverted research microscope Nikon Eclipse Ti-E (Nikon) with the perfect focus system (PFS) (Nikon), equipped with a Nikon CFI Apo TIRF 100×1.49-NA oil objective (Nikon) and a Photometrics Evolve 512 EMCCD (Roper Scientific) camera, and controlled with MetaMorph 7.7 software (Molecular Devices). Images were projected onto the chip of an Evolve 512 camera with an intermediate 2.5× lens (Nikon C mount adaptor 2.5×). To keep in vitro samples at 30 °C, we used an INUBG2E-ZILCS (Tokai Hit) stage-top incubator.

For excitation, we used 491 nm/100 mW Stradus (Vortran) and 561 nm/100 mW Jive (Cobolt) lasers. For simultaneous imaging of green and red fluorescence, we used a triple-band TIRF polychroic filter (ZT405/488/561rpc, Chroma) and triple-band laser emission filter (ZET405/488/561m, Chroma), mounted in the metal cube (91032, Chroma) together with an Optosplit III beam splitter (Cairn Research) equipped with a double-emission-filter cube configured with ET525/50m, ET630/75m and T585LPXR (Chroma) filters.

Doubly cycled GMPCPP-stabilized MT seeds were prepared as described before^50^, by incubating the tubulin mix containing 70% unlabeled porcine brain tubulin (Cytoskeleton), 18% biotin-tubulin (Cytoskeleton) and 12% rhodamine-tubulin (Cytoskeleton) at a total final tubulin concentration of 20 µM with 1 mM GMPCPP (Jena Biosciences) at 37°C for 30 minutes. MTs were pelleted by centrifugation in an Airfuge for 5 minutes at 119,000 × g and then depolymerized on ice for 20 minutes. This was followed by a second round of polymerization at 37°C with 1 mM GMPCPP. MT seeds were then pelleted as above and diluted 10-fold in MRB80 buffer (80 mM PIPES, pH 6.8, supplemented with 4 mM MgCl_2_ and 1 mM EGTA) containing 10% glycerol, snap frozen in liquid nitrogen and stored at −80°C.

The in vitro reconstitution assays with dynamic MTs were performed under the same conditions as described previously^2^. Briefly, after coverslips were functionalized by sequential incubation with 0.2 mg/ml PLL-PEG-biotin (Susos) and 1 mg/ml neutravidin (Invitrogen) in MRB80 buffer, GMPCPP-stabilized microtubule seeds were attached to the coverslips through biotin-neutravidin interactions. Flow chambers were further blocked with 1 mg/ml κ-casein. The reaction mix with purified proteins in MRB80 buffer supplemented with 20 μM porcine brain tubulin, 0.5 μM X-rhodamine-tubulin, 75 mM KCl, 1 mM GTP, 0.2 mg/ml κ-casein, 0.1% methylcellulose and oxygen scavenger mix (50 mM glucose, 400 μg/ml glucose oxidase, 200 μg/ml catalase and 4 mM DTT) was added to the flow chamber after centrifugation in an Airfuge for 5 minutes at 119,000 × g. The flow chamber was sealed with vacuum grease, and dynamic MTs were imaged immediately at 30 °C with a TIRF microscope. All tubulin products for TIRF microscopy were from Cytoskeleton.

### Protein expression and purification for Cryo-EM

Human CAMSAP1 residues 1474-1613 encompassing the CKK domain (HsCKK) were cloned into pET28a vector and expressed in BL21(DE3) cells (Stratagene). Following purification via immobilized metal-affinity chromatography (IMAC) using Ni-NTA resin (Qiagen), the protein was further purified on an ion exchange column MonoS and gel filtration column Superose 6 (GE Healthcare). Purified protein was concentrated to ∼20 mg/ml in BRB20 buffer (20mM PIPES, 2mM MgCl_2_, 1mM EGTA, 1mM DTT, pH 6.8), snap-frozen and stored at −80 °C.

The DNA encoding for the CKK domain of *N. gruberi* CAMSAP (NgCKK, residues 621-788; Uniprot Gene: NAEGRDRAFT_50049) was cloned into the pET-based bacterial expression vector PSPCm2, which encodes for an N-terminal 6x His-tag and a PreScission cleavage site using a positive selection cloning approach^51^. Following protein expression in BL21 (DE3) RIPL cells (Agilent), protein was purified by immobilized metal-affinity chromatography (IMAC) followed by size exclusion chromatography. Purified protein was concentrated to ∼24 mg/ml in BRB20 buffer, snap-frozen and stored at −80 °C. Protein quality and identity were analyzed by SDS-PAGE and mass spectrometry, respectively.

tsA201 cell tubulin was purified from tsA201 cell cultures as described previously^28,52,53^. Briefly, tubulin was isolated from cell lysates via immobilized TOG1 affinity, then tubulin eluted with 0.5M ammonium sulfate. Tubulin was then buffer exchanged into BRB80 buffer (80mM PIPES, 2mM MgCl_2_, 1mM EGTA, 1mM DTT, pH 6.8) with 10% glycerol, and 20 μM GTP and flash frozen in liquid nitrogen. The tubulin was further purified by cycling^54^ then buffer exchanged into BRB80 with 20μM GTP and flash frozen in liquid nitrogen.

### Cryo-EM Sample Preparation

MTs were polymerised using using 5mg/ml tsA201 cell tubulin at 37°C for 45 minutes in BRB80 containing 1 mM GTP. 1mM paclitaxel in DMSO was then added and MTs incubated at 37°C for another 45 minutes. Stabilised MTs were left at room temperature for at least 24 hours then diluted in BRB20 to 0.5mg/ml before use. 4μl of MTs in BRB20 were pre-incubated on glow-discharged holey C-flat^TM^ carbon EM grids (Protochips, Morrisville, NC) at room temperature for 90 seconds, excess buffer manually blotted away, then 4μl of 1mg/ml HsCKK domain or NgCKK added for 45 seconds. Excess buffer was again manually blotted away, followed by a final 4μl application of either HsCKK or NgCKK at the same concentration. Grids were then placed in a Vitrobot Mark IV (FEI Co., Hillsboro, OR) at room temperature and 80% humidity, incubated for a further 45 seconds, then blotted and vitrified in liquid ethane.

### Cryo-EM data collection and processing

Low dose movies were collected manually on a K2 direct electron detector (Gatan) installed on a FEI Tecnai G2 Polara operating at 300kV with a quantum post-column energy-filter (Gatan), operated in zero-loss imaging mode with a 20-eV energy-selecting slit. A defocus range of 0.5-3.5μm and a calibrated final sampling of 1.39Å/pixel was used with the K2 operating in counting mode at 5e-/pixel/second. The total exposure was ∼42e-/Å2 over 16 seconds at 4 frames/sec. Movie frames were aligned using Motioncorr2^55^ with a patch size of 5 to generate full dose and dose-weighted sums. Full dose sums were used for CTF determination in gCTF^56^, then dose-weighted sums used in particle picking, processing and generation of the final reconstructions.

MTs were boxed manually in Relion’s (v3.0) helical mode^25-27^ with a box separation distance of 82Å (roughly the MT dimer repeat distance) and further processed using a custom pipeline designed to account for the pseudo-helical nature of MTs with a seam. The pipeline, inspired by previous pipelines^57,58^ developed for Spider/Frealign^59,60^, is based in RELION and uses accessory scripts to place refinement and classification restraints on individual MTs using prior knowledge of MT geometry. Briefly, the protofilament number of all segments within each MT was assigned according to the modal class of those segments after one iteration of 3D classification to low-pass filtered references of 11-16-protofilament MTs. Only the dominant 13-protofilament and 14-protofilament classes were separately processed further. Rough alignment parameters of each MT to its corresponding low-pass filtered 13-protofilament or 14-protofilament reference were assigned. On the basis of ϕ angles determined for each segment, median ϕ angles were assigned to all segments in a given MT. Assigned ϕ angles for each MT were checked by 3D classification against low-pass filtered MT references rotated and translated to represent all possible seam positions and αβ-tubulin registers (i.e 26 references for a 13-protofilament MT, with 13 seam positions and their counterparts translated 1 monomer along the helical axis). Rough final ϕ angles were assigned according to the modal 3D class of all segments within each MT. Fine local refinement was then performed, without applied helical symmetry. Subsequently, 1 iteration each of Bayesian polishing and per-particle CTF refinement was performed in Relion v3.0^25^, followed by a final round of fine local refinement with or without applied helical symmetry.

4 × binned data was used in all processing steps except the final 3D refinement (1 × binned data) and segment averages of 7 segments along the helical axis were used for 3D classification steps. Final global resolutions are estimated from the Fourier shell correlation 0.143 cut-off of gold-standard FSCs generated by applying soft masks to the central 15% portion of the two independent half-maps along the helical axis. Reconstructions used for model building, refinement and display were sharpened (using globally determined B-factors, see Supplementary Table 1) to local resolution cut-offs determined using Relion v3.0’s internal local resolution program (see Supplementary Figure 1).

### Cryo-EM Model Building and Refinement

α1B/βI+βIVb tsA201 cell tubulin with bound HsCKK or NgCKK was modelled into pseudo-symmetrised density maps via iterative rounds of direct model building in Coot^61^ and real-space refinement applied in Phenix^62^. Tubulin in the starting models was constructed by fitting dimers from the GDP-EB3 13pf microtubule cryo-EM structure (PDB 3JAR^41^), into corresponding densities and applying mutations to account for sequence differences and incorporating taxol from the structure of tubulin-taxol zinc sheets (PDB 1JFF^63^). NgCKK and HsCKK domain starting models were created via homology modelling of the x-ray and NMR structures of CAMSAP3 CKK domain (PDBs 5LZN^2^ and 1UGJ (unpublished)) using Modeller^64^. Starting models of HsCKK or NgCKK domains were then fitted into density alongside tubulin and merged into single starting models constructed of 6 tubulin dimers and 4 CKK domains. Cryo-EM figures were prepared using UCSF Chimera^65^.

### Protein preparations for NMR

Human CAMSAP1 N1492A CKK (residues 1474-1613) was cloned into a pET28a vector. For sample preparation of low MAS ssNMR, uniformly [^13^C, ^15^N]-labeled CAMSAP1 N1492A CKK was produced in *E. coli* strain Rosetta 2 in M9 minimum medium supplemented with 25 μg/ml kanamycin and 35 μg/ml chloramphenicol. Cells were induced when the OD600 reached 0.6 with 0.3 mM IPTG at 25 °C for 5 hrs. For ^1^H detected ssNMR experiments, uniformly [^2^H, ^13^C, ^15^N]-labeled CKK mutant was produced in *E. coli* Rosetta 2 strain in M9 minimum medium that was prepared with D_2_O, deuterated 13C-glucose and 15N-NH4Cl. When OD600 reached 0.6, 0.3 mM IPTG was added for induction at 25 °C for 5 hrs. The proteins were purified by a ÄKTA pure system with a POROS™ MC column that was saturated with Ni^2+^. The column was first equilibrated with washing buffer (50 mM phosphate buffer, pH 8, 200 mM NaCl, 1 mM β-mercaptoethanol and 20 mM imidazole). Proteins were eluted with the same buffer but containing 400 mM imidazole. Proteins were then loaded onto a SEC HiLoad Superdex 75 26/60 column (GE Healthcare) that was equilibrated with 40 mM phosphate buffer, pH 7, supplemented with 150 mM NaCl and 1 mM DTT. Subsequently, the labeled proteins were concentrated and used for ssNMR sample preparation.

To prepare CKK-MT complexes for low-speed MAS ssNMR, 20 mg of lyophilized porcine tubulin was dissolved in BRB80 buffer to a final concentration of 2 mg/ml. Microtubule polymerization was done with 1 mM GTP and addition of 20 μM paclitaxel (Sigma) for 30 min at 30 °C. Paclitaxel-stabilized MTs were pelleted down at 180,000 × g (Beckman TLA-55 rotor) at 30 °C for 30 min and resuspended in warm BRB80 buffer with 20 μM paclitaxel. [^13^C, ^15^N]-labeled CKK N1492A was then added to a final concentration of 65.3 μM (4:1 CKK/tubulin) and incubated at 37 °C for 30 min. The pellet was centrifuged down at 180,000 × g (Beckman TLA-55 rotor) at 30 °C for 30 min and washed with 40 mM phosphate buffer, pH 7, without disturbing the pellet. The pellet was then transferred and packed into a 3.2 mm rotor.

To prepare the CKK-MTs complexes for ^1^H detected experiments, uniformly [^2^H, ^13^C, ^15^N]-labeled CKK was first purified and maintained in the protonated buffer overnight to allow back-exchange of amide protons. 5 mg lyophilized porcine brain tubulin was dissolved in 2.5 mL BRB80 buffer to make a concentration of 2 mg/mL. Tubulin was then polymerized with 1 mM GTP and 20 μM paclitaxel for 30 min at 30 °C. Paclitaxel-stabilized MTs were then ultracentrifuged at 180,000 × g at 30°C for 30 min and then resuspended with warm BRB80 buffer with 20 μM paclitaxel. The CKK domain was added to the resuspended MTs and incubated at 37°C for 30 min. The CKK-MTs complexes were then ultracentrifuged at 180,000 × g at 30°C for 30 min. Finally, the pellet was washed with phosphate buffer and packed into a 1.3 mm NMR rotor.

For sample preparation for solution-state NMR, uniformly [^13^C, ^15^N]-labeled and ^15^N-labeled CAMSAP1 N1492A CKK were expressed and purified with the same way as described above and supplemented with 5% D_2_O for solution-state NMR measurements.

### NMR experiments and data analysis

Resonance assignments of CAMSAP1 CKK N1492A were obtained from standard solution-state NMR experiments (2D HSQCs, 3D HNCA, HNCO, HNCACB, CBCA(CO)NH) on free [^13^C, ^15^N]-labeled CKK recorded on a 600 MHz spectroscopy (Bruker Biospin). Standard MAS ssNMR experiments were conducted on a 950 MHz standard-bore spectrometer (Bruker Biospin) equipped with a 3.2 mm triple-channel MAS HCN probe. The experiments include 2D ^13^C-^13^C proton-driven spin-diffusion (PDSD)^66,67^ and NCA experiments ^68^ (set temperature 260 K, MAS rate 14 kHz). The spin diffusion mixing time was set to 30 ms, and a SPECIFIC-CP^69^ transfer time of 2.2 ms was employed for the NCA experiment. Fast MAS, ^1^H detected, ssNMR experiments were performed on a 800 MHz wide-bore spectrometer (Bruker Biospin) equipped with a 1.3 mm triple-channel MAS HXY probe. The experiments included 2D NH and 3D CANH^70^ experiments (set temperature 244 K, MAS rate 55 kHz). ssNMR MT samples were stable over time as confirmed by comparing ssNMR spectra at standard and fast MAS and by conducting negative staining EM experiments before and after ssNMR measurements.

CPMG relaxation dispersion^31^ and CEST measurements^32^ were based on the 2D ^1^H-^15^N HSQC spectra and were recorded as pseudo 3D on the ^15^N-labeled CAMSAP1 N1492A CKK. The acquisition times in each 2D plane are 66 ms for ^1^H (direct dimension) and 48.6 ms for ^15^N (indirect dimension). CPMG relaxation dispersion experiments were conducted with temperature compensation and single scan interleaved. The data were measured at CPMG fields of 50, 150, 250, 350, 450, 550, 650, 750, 850, 950, 1050, 1150, 1250, 1400 and 1500 Hz, which all applied for a constant transverse relaxation time of 40 ms. The saturation during CEST experiments were carried out with a 400 ms pulse of 15 Hz radio frequency field strength on ^15^N. The saturation offsets ranged between 8200 and 6325Hz with a spacing of 25 Hz on ^15^N.

The difference of chemical-shift values between the free- and bound-state CKK were first translated into the units of linewidths in corresponding dimensions. Then the differences in ^1^H and ^15^N dimensions were combined as 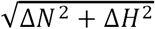 The signal linewidth in ^1^H and ^15^N dimensions were determined to amount to 0.1 and 0.6 ppm, respectively.

CMPG data were processed as follows: For each residue, the standard deviation and the average of signal intensities with different CPMG frequencies were calculated. The ratio of these two values was plotted for every residue on the structures.

For determining chemical-shift changes between free and MT bound CAMSAP1 CKK and CAMSAP1 CKK N1492A, we transferred solution-state NMR shifts obtained on the free variants to ssNMR experiments on the complexes assuming spectral proximity in all three independent dimensions (HN, N, Cα). With this strategy, we were able to transfer 48 backbone assignments as demonstrated in Supplementary Figure 5c.

### Data Availability

The 13-protofilament and 14-protofilament HsCKK and NgCKK-MT models along with their corresponding electron density maps are deposited in the PDB. The PDB codes are as follows: 13-protofilament HsCKK-MT, PDB: 6QUS, 14-protofilament HsCKK-MT, PDB: 6QVJ, 13-protofilament NgCKK-MT, PDB: 6QUY, 14-protofilament NgCKK-MT, PDB: 6QVE. The EMDB codes (C1 reconstruction and symmetrised asymmetric unit) are as follows: 13-protofilament HsCKK-MT, EMDB-4643, 14-protofilament HsCKK-MT, EMDB-4654, 13-protofilament NgCKK-MT, EMDB-4644,14-protofilament NgCKK-MT, EMDB-4650. All supporting data are available from the authors on request, and/or are available in the manuscript itself.

## SUPPLEMENTARY FIGURES

**Supplementary Table 1.**
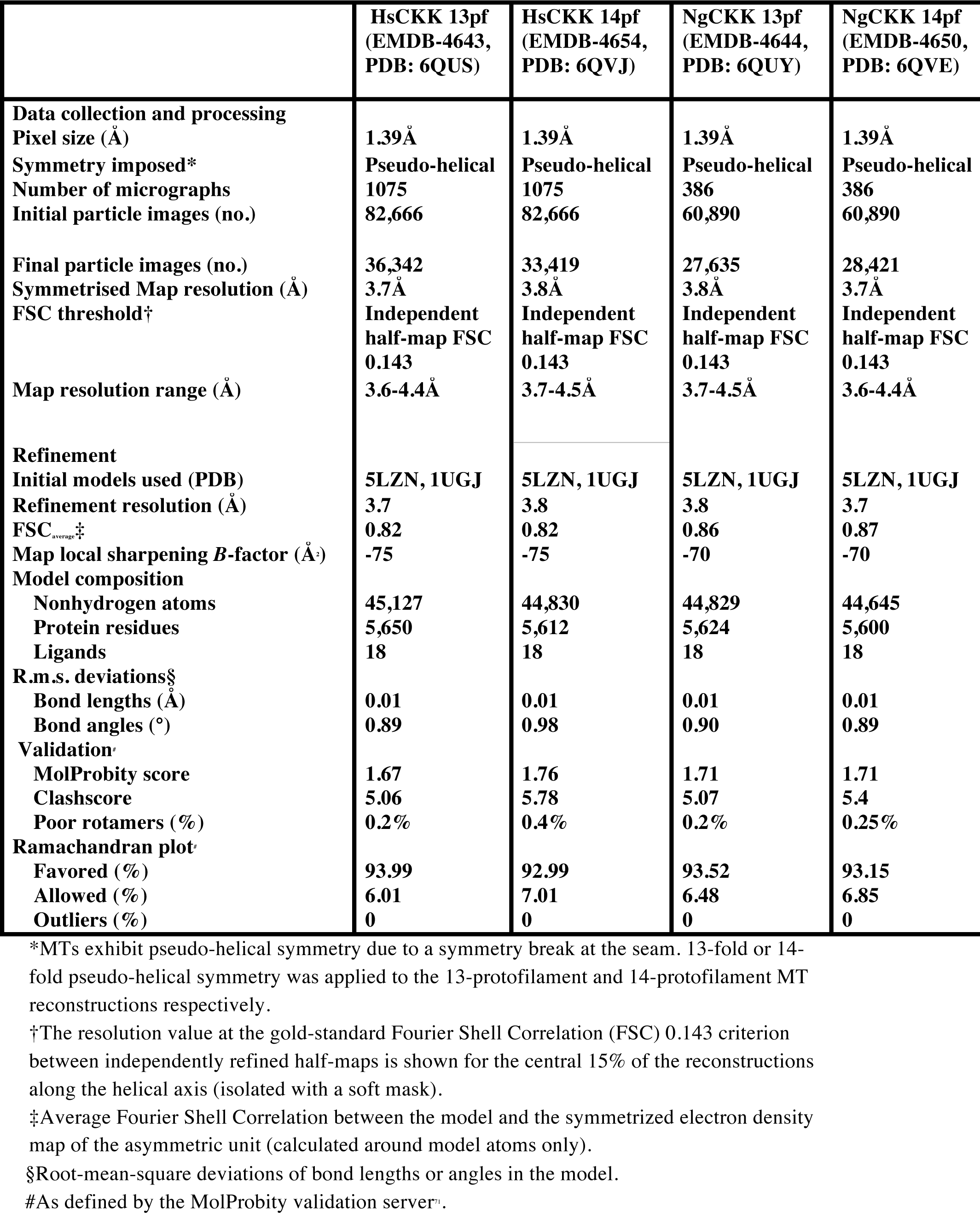
Cryo-EM reconstruction information and model refinement statistics and model geometry. Data collection and processing information for the HsCKK and NgCKK 13-protofilament and 14-protofilament MT datasets and model refinement statistics for the tubulin + CKK asymmetric unit models built into their respective the symmetrized electron density maps.

**Supplementary Table 2.** Helical parameters for CKK-MT reconstructions. 3-start rise and twist were calculated directly from the reconstructions in RELION v3.0’s helical mode^25-27^. Intradimer, interdimer and dimer repeat distances were calculated directly from the refined models in Chimera^65^. Dimer Skew was measured as azimuthal (ϕ) angle change per dimer in the aligned data sets as described for Figure 4b.

|  | **3-start**<br>**rise (Å)** | **3-start**<br>**twist (°)** | **Intradimer**<br>**distance (Å)** | **Interdimer**<br>**distance (Å)** | **Dimer**<br>**distance (Å)** | **Dimer**<br>**Skew (°)** |
| --- | --- | --- | --- | --- | --- | --- |
| **CAMSAP1-**<br>**CKK 13pf-**<br>**taxol** | 9.49 | -27.65 | 41.3 | 41.00 | 82.3 | 0.50 |
| **N.Grub-**<br>**CKK 13pf-**<br>**taxol** | 9.47 | -27.67 | 41.1 | 40.8 | 82.0 | 0.14 |
| **CAMSAP1-**<br>**CKK 14pf-**<br>**taxol** | 8.80 | -25.72 | 41.2 | 40.8 | 82.0 | -0.04 |
| **N.Grub-**<br>**CKK 14pf-**<br>**taxol** | 8.80 | -25.72 | 41.1 | 40.9 | 82.0 | -0.28 |

**Supplementary Figure 1.**
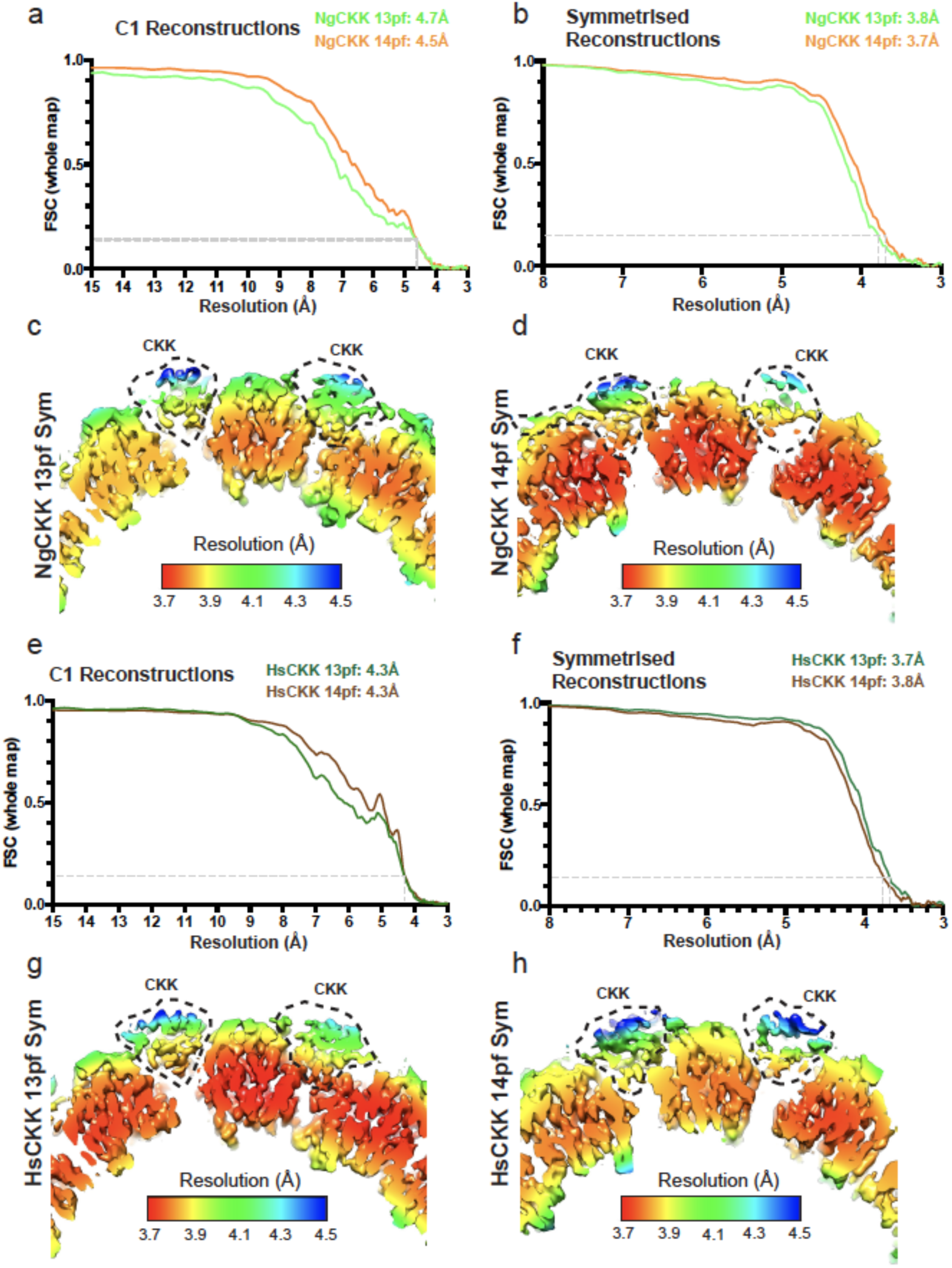
Evaluation of the resolutions of the CKK-MT reconstructions. a) FSC curves for the NgCKK-MT 13-(green) and 14-protofilament (orange) C1 reconstructions, with the 0.143 cut-off indicated; b) FSC curves for the NgCKK-MT 13-(green) and 14-protofilament (orange) symmetrised reconstructions, with the 0.143 cut-off indicated; c) Local resolution depiction using RELION v3.0’s local resolution determination of a slice through the NgCKK-MT symmetrised 13-protofilament reconstruction viewed from the MT minus end; d) Local resolution depiction using RELION v3.0’s local resolution determination of a slice through the NgCKK-MT symmetrised 14-protofilament reconstruction viewed from the MT minus end; e) FSC curves for the HsCKK-MT 13-(green) and 14-protofilament (brown) C1 reconstructions, with the 0.143 cut-off indicated; f) FSC curves for the HsCKK-MT 13-(green) and 14-protofilament (brown) symmetrised reconstructions, with the 0.143 cut-off indicated; g) Local resolution depiction using RELION v3.0’s local resolution determination of a slice through the HsCKK-MT symmetrised 13-protofilament reconstruction viewed from the MT minus end; h) Local resolution depiction using RELION v3.0’s local resolution determination of a slice through the HsCKK-MT symmetrised 14-protofilament reconstruction viewed from the MT minus end;

**Supplementary Figure 2.**
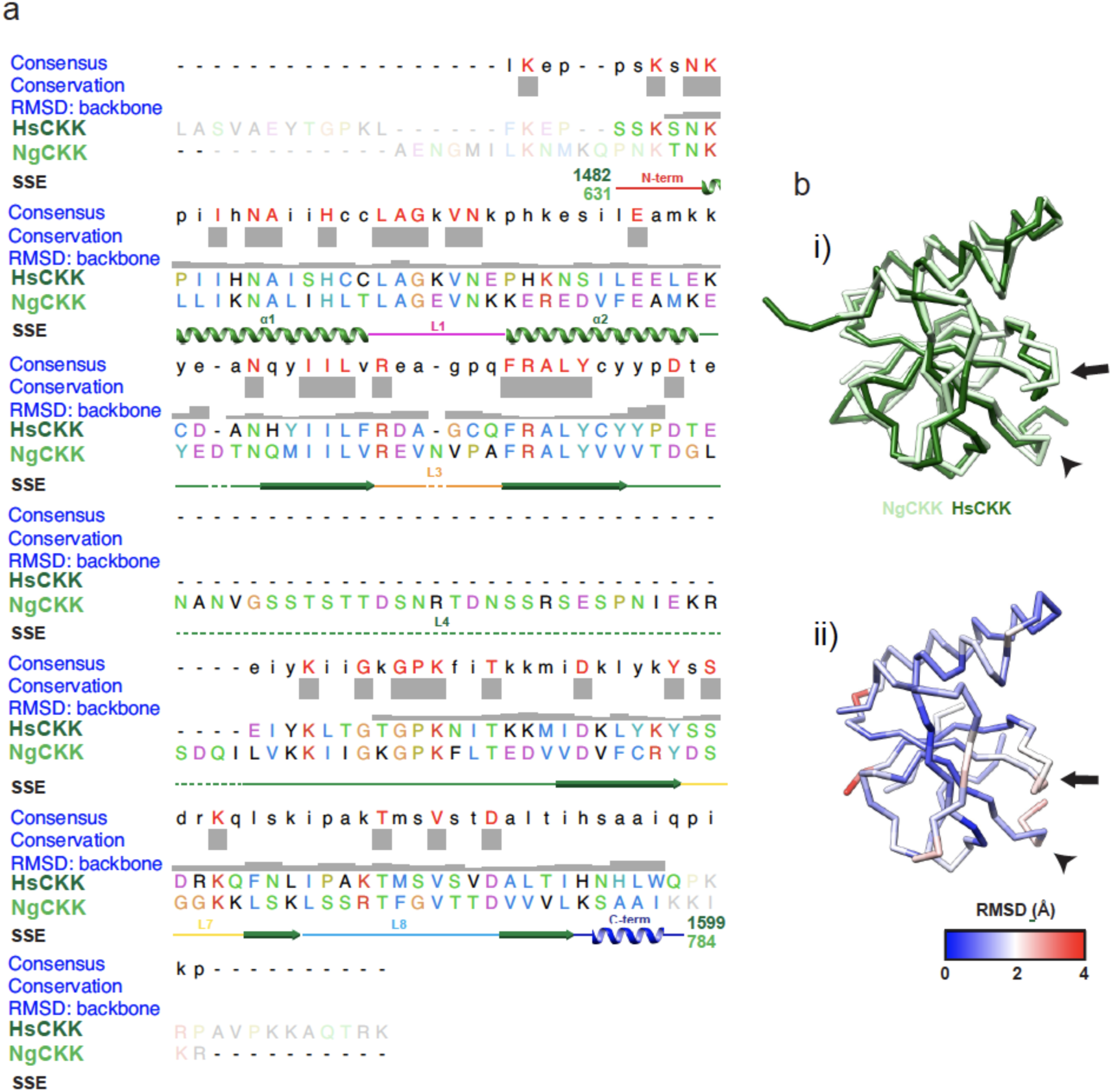
Comparison of NgCKK and HsCKK primary and 3D structures. a) Sequence alignment of HsCKK and NgCKK (extracted from our previous larger CKK domain sequence alignment^2^) annotated in UCSF Chimera^65^ with sequence consensus, conservation, backbone RMSD and secondary structure elements. The N-and C-terminal residue number for the CKK domain within each full-length protein are also indicated; b) i) Overlay of aligned NgCKK (light green) and HsCKK (dark green) backbone models viewed from the MT lumen; ii) Backbone RMSD between NgCKK and HsCKK shown on the NgCKK structure. In both panels, the arrow indicates loop3 and the arrowhead indicates the CKK C-terminus, regions of notable structural divergence between NgCKK and HsCKK.

**Supplementary Figure 3.**
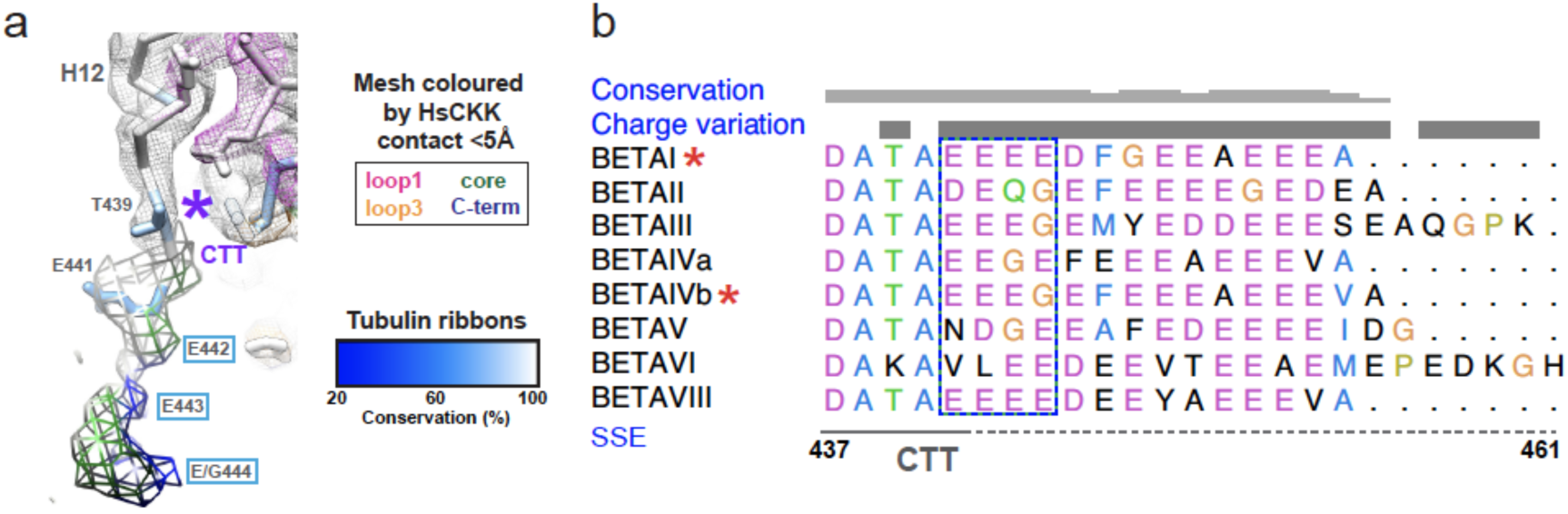
Visualisation of β-tubulin’s CTT interaction with HsCKK is likely facilitated by the homogeneity of tsA201 cell tubulin. a) Density isolated from the HsCKK-MT cryo-EM reconstruction, with the region corresponding to the ordered β-tubulin CTT shown as thick mesh; the sequence of the well-ordered CTT region in tsA201-cell tubulin is shown with unmodelled residues labelled in light blue boxes; the mesh is coloured according to the closest secondary structural element of HsCKK (see key); b) Human β-tubulin isoform sequences (letters coloured in Clustal-X style by residue subtype) for C-terminal tail residues 438-444, showing that the BETAI and BETAIVb isoforms present in tsA201 cell tubulin (indicated by a red *) only differ in their final residue. Blue boxed region indicates residues that were not fully modelled, although density corresponding to this region is visualized and forms contacts with both the CKK core and C-terminus.

**Supplementary Figure 4.**
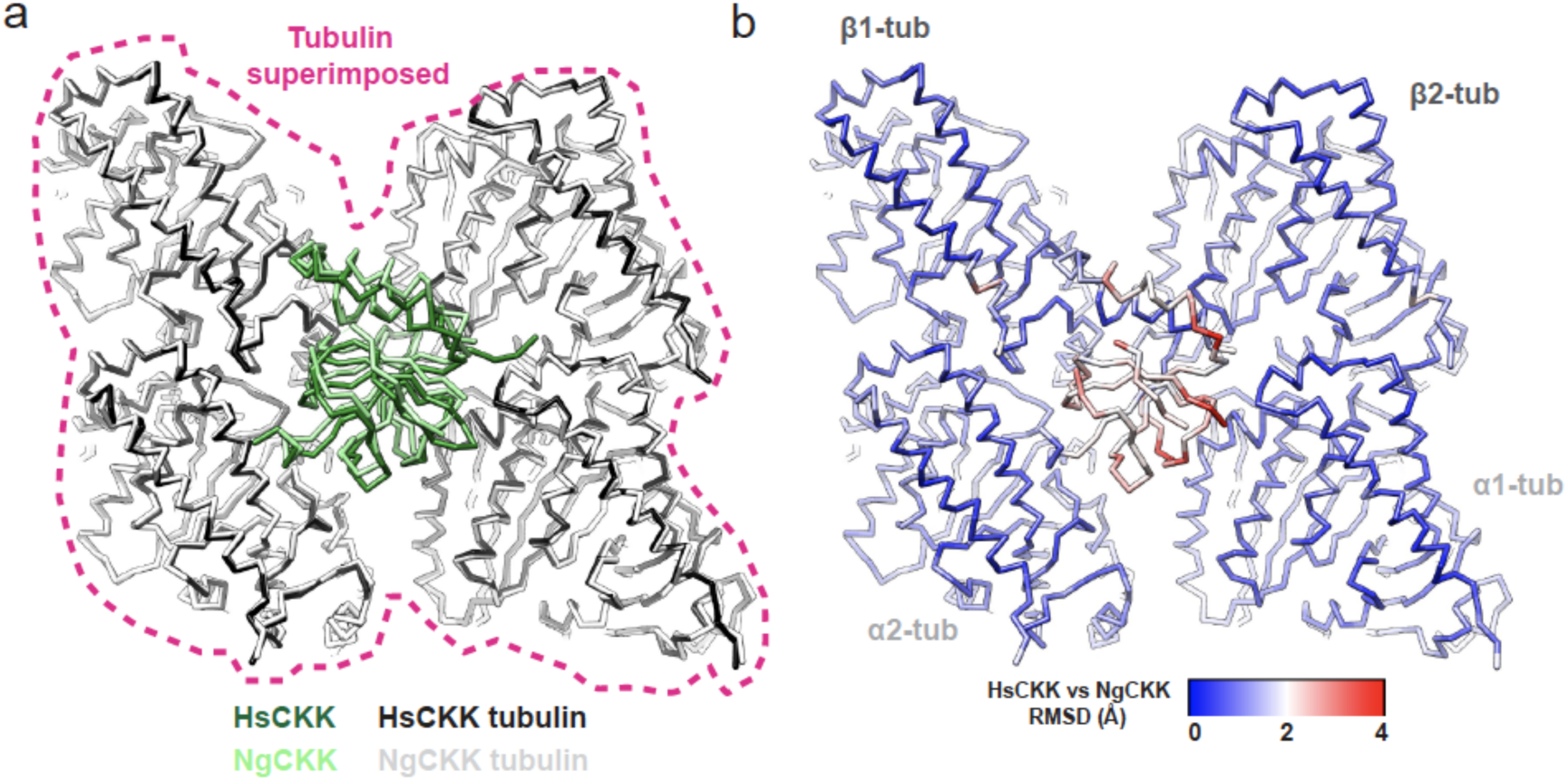
Structural differences in MT interaction between NgCKK and HsCKK. a) Backbone models of tubulin dimer pairs from the NgCKK-MT (light grey) and HsCKK-MT (dark grey) structures align very well, while the CKK domains (NgCKK, light green; HsCKK, dark green) exhibit differences especially adjacent to the α-tubulins (annotated in detail in Fig. 2a); b) Backbone RMSD between these models shown on the NgCKK-MT model; models are viewed from the outside MT surface in a) and b).

**Supplementary Figure 5.**
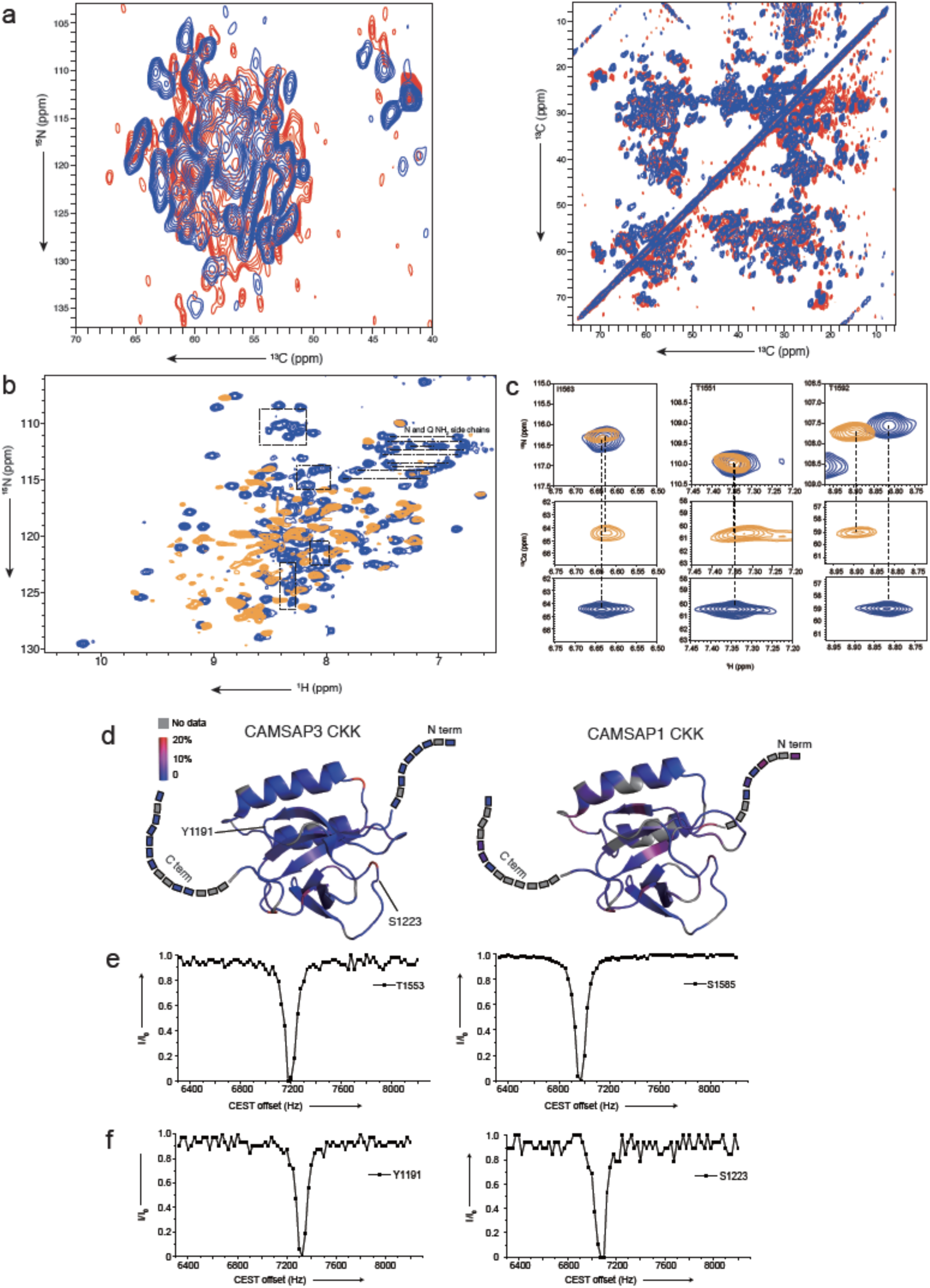
Rigidity of the HsCKK domain revealed by high-resolution NMR. a) Comparison of solid-state NMR spectra obtained on WT (red, CAMSAP3 CKK) and mutant (blue, CAMSAP1 CKK N1492A) CKK. Data are shown from (i) 2D NCA and ii) 2D PDSD spectra; b) NMR Chemical-shift perturbations of CAMSAP1 CKK N1492A upon binding to MTs. 2D NH correlations are shown for free CAMSAP1 CKK N1492A in solution (blue) and MT bound CKK (orange). For clarification, NH2 correlations from the side chains of N and Q residues are indicated by dashed lines. Furthermore, signals from purification tags are highlighted (boxes) that do not appear in the ssNMR NH correlation spectrum (orange) of the MT-bound CKK; c) Spectral cutouts comparing solid-state NMR signal sets of free (blue) and MT bound (orange) HsCKK_N1492A for three selected residues. Top panels show the zoom-in regions from the 2D ^15^N-^1^H correlated spectra, while the middle and bottom panels reflect regions from 3D CANH (solid state) and HNCA (liquid state) spectra at corresponding ^15^N chemical shifts, respectively. See panel b for full spectra; d) Protein Dynamic-sensitive NMR profiles for wildtype CKK domains. Comparison of wild type (i) CAMSAP3 CKK and (ii) CAMSAP1 CKK transverse relaxation rates with same CPMG setting as Fig. 5aii; e) Solution CEST profiles of free CKK shown for residues T1553 and S1585, which showed significant chemical shift perturbation upon binding to MTs (see Fig. 5ai); f) CEST profiles of Y1191 and S1223 from CAMSAP3 CKK exhibiting similar dynamics profiles as shown for T1553 and S1585 from CAMSAP1 CKK N1492A panel e.

**Supplementary Figure 6.**
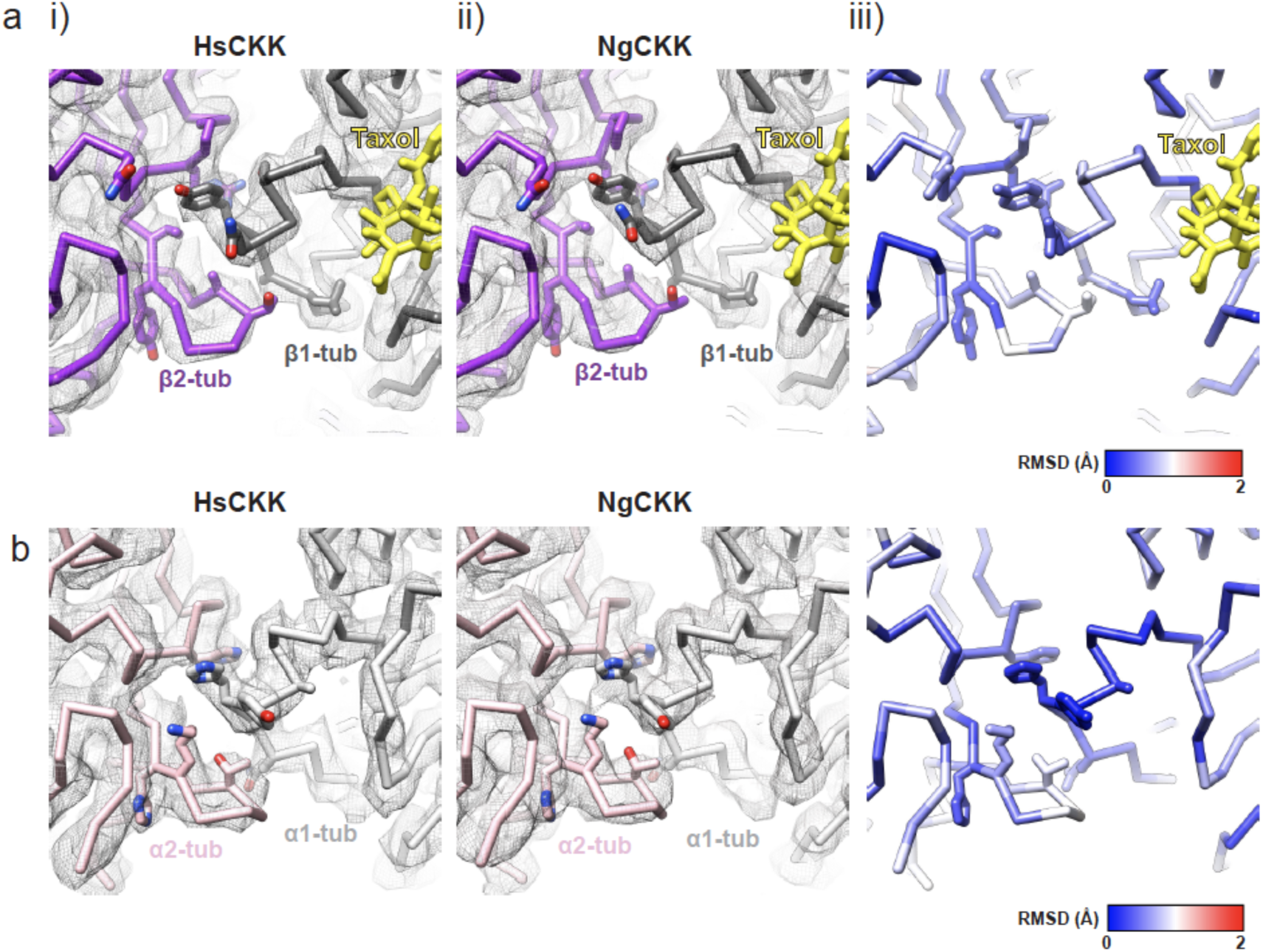
HsCKK MT remodelling does not cause detectable differences in the inter-protofilament lateral contacts. a) Comparison of the β-tubulin lateral contacts viewed from the MT lumen in the 13-protofilament i) HsCKK reconstruction and model, ii) NgCKK reconstruction and model iii) backbone RMSD of this region, showing that the remodelling of the MT architecture does not cause detectable difference in the β-tubulin interprotofilament lateral contacts; β1-tubulin is coloured dark grey, β2-tubulin is coloured purple and taxol is shown in yellow; b) Comparison of the α-tubulin lateral contacts viewed from the MT lumen in the 13-protofilament i) HsCKK reconstruction and model, ii) NgCKK reconstruction and model iii) backbone RMSD of this region, showing that the remodelling of the MT architecture does not cause detectable difference in the α-tubulin interprotofilament lateral contacts; α1-tubulin is coloured light grey and α2-tubulin is coloured pink.

## Supplementary Movies

**Supplementary Movie 1. Comparison of NgCKK and HsCKK binding positions on two laterally adjacent tubulin dimers.** The pair of neighbouring tubulin dimers in 13-protofilament HsCKK and NgCKK models were superimposed and a morph created between the two, demonstrating structural and positional differences between the HsCKK and NgCKK relative to the MT lattice. The CKK models are coloured according to SSE/loop colour scheme in Fig 1.

**Supplementary Movie 2. NgCKK is minimally displaced from its binding site in 14-compared to 13-protofilament MTs.** A transverse view from the MT plus end through 3 protofilaments from the NgCKK-MT 13-morphing to 14-protofilament models (backbone representation) superimposed on the central protofilament. This shows how the adjacent protofilaments adopt a shallower relative angle in 14-protofilament MTs and reveals the minimal displacement of bound NgCKK in response to this change in lateral protofilament curvature. NgCKK is shown in light green, β-tubulin in dark grey, α-tubulin in light grey.

**Supplementary Movie 3. HsCKK is displaced from its binding site in 14-compared to 13-protofilament MTs, demonstrating its greater sensitivity to microtubule lateral curvature.** A transverse view from the MT plus end through 3 protofilaments from the HsCKK-MT 13-morphing to 14-protofilament models (backbone representation) superimposed on the central protofilament. This shows how the adjacent protofilaments adopt a shallower relative angle in 14-protofilament MTs and reveals the displacement of bound HsCKK in response to this change in lateral protofilament curvature. HsCKK is shown in green, β-tubulin in dark grey, α-tubulin in light grey.

